# Chemical induction of gut β-like-cells by combined FoxO1/Notch inhibition as a glucose-lowering treatment for diabetes

**DOI:** 10.1101/2021.12.07.471572

**Authors:** Takumi Kitamoto, Yun-Kyoung Lee, Nishat Sultana, Wendy M. McKimpson, Hitoshi Watanabe, Wen Du, Jason Fan, Bryan Diaz, Hua V. Lin, Rudolph L. Leibel, Sandro Belvedere, Domenico Accili

**Affiliations:** Department of MedicineVagelos College of Physicians and Surgeons, Columbia University, New York, NY 10032; Naomi Berrie Diabetes Center Vagelos College of Physicians and Surgeons, Columbia University, New York, NY 10032; Forkhead BioTherapeutics, Inc., New York, NY; Department of Pediatrics Vagelos College of Physicians and Surgeons, Columbia University, New York, NY 10032; BioFront Therapeutics, Beijing, China; Department of Ophthalmology, Bascom Palmer Eye Institute, University of Miami, Miami, FL, 33146; Chiba University Graduate School of Medicine, Chiba, Japan, 2608670

**Author notes:** Corresponding Author: Takumi Kitamoto, MD, PhD, Department of Medicine, Vagelos College of Physicians and Surgeons, Columbia, University, New York, NY. Y.K.L and N.S. contributed equally. Reprint requests: Domenico Accili.

**Keywords:** Diabetes, Insulin, β cell replacement, FOXO1, FoxO1 inhibitor, Notch inhibition

## Abstract

Lifelong insulin replacement remains the mainstay of type 1 diabetes treatment. Genetic FoxO1 ablation promotes enteroendocrine cell (EECs) conversion into glucose-responsive β-like cells. Here, we tested whether chemical FoxO1 inhibitors can generate β-like gut cells. Pan-intestinal epithelial FoxO1 ablation expanded the EEC pool, induced β-like cells, and improved glucose tolerance in *Ins2*^Akita/+^ mice. This genetic effect was phenocopied by small molecule FoxO1 inhibitor, Cpd10. Cpd10 induced β-like cells that released insulin in response to glucose in mouse gut organoids, and this effect was strengthened by the Notch inhibitor, DBZ. In *Ins2*^Akita/+^ mice, a five-day course of either Cpd10 or DBZ induced insulin-immunoreactive β-like cells in the gut, lowered glycemia, and increased plasma insulin levels without apparent adverse effects. These results provide proof of principle of gut cell conversion into β-like cells by a small molecule FoxO1 inhibitor, paving the way for clinical applications.

**One Sentence Summary:** Orally available small molecule FoxO1 inhibitor phenocopied genetic FoxO1 ablation in generating gut β-like cells

## INTRODUCTION

Efforts to generate insulin-producing cells that can be delivered to patients in vivo to treat type 1 diabetes have been underway for decades. Decisive progress has been made in the generation of pancreatic hormone-producing cells from embryonic stem (ES) or induced pluripotent stem (iPS) cells ^1^. These strategies can provide an unlimited source of β-cells to address the scarcity of islet donors. However, there are at least two significant hurdles for cell-based replacement therapies: physical limitations in delivering sufficient quantities of cells, and protection from autoimmunity to maintain their function ^2^.

An alternative approach entails conversion of endogenous non-β cells into insulin-secreting cells ^3^. However, this approach suffers from poor reproducibility of claims. We and others have shown that gut epithelial cells can be reprogrammed into functional glucose-sensing β-like cells ^4–8^. Genetic ablation of FoxO1 in EECs in vivo gives rise to β-like cells that produce and secrete insulin like endogenous β-cells ^6^. In addition, FoxO1 inhibition by shRNA or a dominant-negative mutant converts cells in human iPS cell-derived gut organoids to pancreatic β-like cells^7^, suggesting that the ability of EECs to become insulin-producing cells is a shared property of rodent and human gut cells. This approach provides an attractive alternative strategy by which to replace β-cell mass in type 1 diabetes. In addition, gut cells can potentially escape autoimmune attack owing to their rapid turnover, scattered distribution, and immune privilege ^9^.

To translate this interesting biology into a treatment, it is necessary to inhibit FoxO1 pharmacologically. As a member of the forkhead DNA binding domain family ^10–12^, FoxO1 lacks a ligand binding domain, complicating the search for an appropriate small molecule agonist or antagonist. However, we have identified at least two classes of highly specific ^13^ and selective FoxO1 inhibitors derived from an unbiased chemical library screen ^14^. In the present study, we asked whether chemical FoxO1 inhibition *in vivo* can phenocopy the effects of somatic ablation of FoxO1 in gut epithelial cells on EEC differentiation. In addition, in light of previous findings ^15, 16^, indicating that the FoxO and Notch pathways cooperate on pancreatic β-cell development and of the effect of Notch on this process ^17^, we also examined the effect of combining inhibitors of the two pathways on EEC differentiation and diabetes treatment. Our findings establish the feasibility of delivering a small molecule FoxO1 inhibitor to generate β-like cells in the gut and lower glycemia in insulin-deficient mice.

## RESULTS

### Gut FoxO1 ablation expands the EEC progenitor pool

To investigate the mechanism by which FoxO1 ablation affects EEC formation, we generated *ACTB-tdTomato-EGFP; Neurog3-Cre:FoxO1^fl/fl^* mice and assessed EEC number, localization, and gene expression patterns. The reporter allele yields green fluorescence after cre recombination and allows to monitor Neurog3 expression by comparing GFP^+^ (Neurog^+^) and RFP^+^ (Neurog^−^) cells. In wild-type animals (ACTB-tdTomato-EGFP; Neurog3-Cre, hereafter TG-WT), EECs were located throughout the villi. Following FoxO1 ablation in EEC progenitors (ACTB-tdTomato-EGFP; Neurog3-Cre (+): FoxO1^fl/fl^, hereafter TG-NFKO), EECs were enriched in crypts (Fig. 1A and B). Both GFP^+^ and RFP^+^ cells were readily identified by fluorescence-activated cell sorting (FACS) (P4 and P5 in Fig. 1C); however, GFP^+^ cells appeared to segregate in two different subsets that differed by the strength of the RFP^+^ signal (P5 and P6 in Fig. 1C). mRNA measurements indicated that the three subsets corresponded to RFP^+^GFP^-^ endocrine progenitors (Lgr5^low^, Ngn3^low^, NeuroD^low^, FoxO1^high^, Tph1^low^, ChgA^low^), GFP^+^RFP^-^ terminally differentiated EECs (Lgr5^low^, Ngn3^low^, NeuroD^high^, FoxO1^low^, Tph1^high^, ChgA^high^) and RFP^+^GFP^+^ cells (Lgr5^high^, Ngn3^high^, NeuroD^high^, FoxO1^medium^, Tph1^medium^, ChgA^medium^) that likely represent different stages of EEC maturation from early (P5) to late progenitors (P6) (Fig. 1D–F, S1A–C).

**Figure 1.**
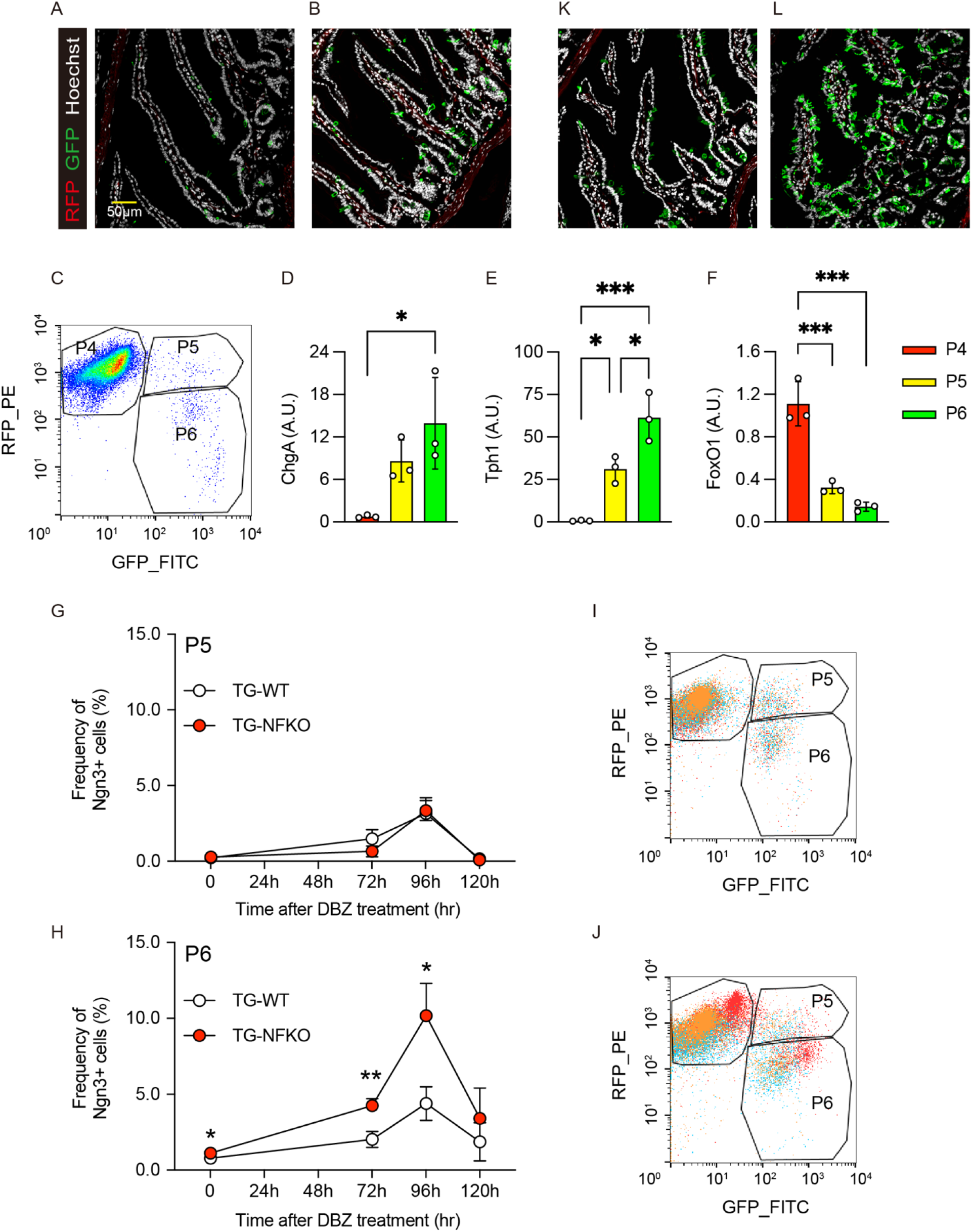
Gut FoxO1 ablation expands the EEC progenitor pool. (A, B) Representative images of GFP and RFP immunofluorescence in intestinal duodenum sections of TG-WT (A) and TG-NFKO (B) mice. (C) Representative FACS plot of TG-WT gut epithelial cells. P4: RFP^+^; P5: RFP^+^/GFP^+^; P6: GFP^+^. (D–F) qPCR of ChgA (D), Tph1 (E), and FoxO1 (F) using mRNA from sorted P4, P5, and P6 cells. N=3 for each group. (G–J) Time course of induction of P5 and P6 population after DBZ treatment. White and red circles indicate TG-WT and TG-NFKO, respectively. 3 × 10^5^ cells were analyzed in each experiment, and each group consists of biological triplicates. Merged FACS plots of three samples from TG-WT (I) and TG-NFKO (J) following 96-hr of DBZ treatment are shown. (K, L) Representative GFP and RFP immunofluorescence images in intestinal duodenum sections of TG-WT (K) and TG-NFKO (L) after 96-hr DBZ treatment. TG-WT: ACTB-tdTomato,-EGFP; Neurog3-Cre: TG-NFKO, ACTB-tdTomato-EGFP; Neurog3-Cre (+): FoxO1^fl/fl^. Data are presented as means ± SD. Scale bar = 50 μm. *= p < 0.05; **= p < 0.01; ***= p < 0.001.

Notch inhibition has been shown to expand the EEC pool ^18^. Thus, we compared the combined effect of FoxO1 ablation and Notch blockade by the γ-secretase inhibitor, DBZ, on the different stages of the EEC lineage (Fig. 1G–J) ^16, 19^. As reported, 96-hr after treatment with a single dose of DBZ, there was a substantial increase of serotonin (5HT), somatostatin (SS), and glucagon-like peptide-1 (GLP-1) cells in C57/BL6 mice (Fig. S2A–C) ^18^. We confirmed that the total number of GFP^+^ EEC was significantly increased at baseline and 96-hr after FoxO1 ablation (Fig. S1D). We next compared the effect of DBZ on FAC-sorted P5 and P6 cells isolated from WT or TG-NFKO mice. In contrast to early progenitors P5 (Fig. 1G), FoxO1 knockout increased the number of late progenitors P6 by 30% (0.79 ± 0.07 vs. 1.12 ± 0.11: *p* = *0.01*) at baseline. DBZ increased EEC number four-fold in TG-WT, and more than eight-fold in TG-NFKO (4.39 ± 1.11 vs. 10.17 ± 2.13: *p* = *0.014*) (Fig. 1G–H). We also assessed EEC localization following 96-hr of DBZ treatment (Fig. 1K-L). In contrast to pre-treatment, the number of GFP^+^ cells in the crypts increased in both control and TG-NFKO mice, while more EECs were detected in the villi of TG-NFKO mice. Thus, Neurog3-driven FoxO1 ablation and Notch inhibition synergistically expanded the EEC progenitor pool and increased EEC number throughout the villi.

### Time course of EEC differentiation after FoxO1 ablation

To test whether the timing of FoxO1 ablation affects the EEC pool, we ablated FoxO1 at the EEC progenitor stage in Neurog3^+^ cells ^20^ or at the epithelial progenitor stage in villin^+^ cells ^21^. We isolated crypts from Neurog3-Cre (+); FoxO1^fl/fl^ (NFKO) or Villin-Cre (+); FoxO1^fl/fl^ (VFKO) mice to prepare primary organoid cultures (mGO) and assessed the time course of FoxO1 and EEC marker expression after DBZ treatment *in vitro*. In basic medium few mGOs were differentiated into EECs. DBZ treatment efficiently differentiated mGOs into EECs, and FoxO1 expression decreased in WT mGO with EEC differentiation (Fig. 2A), while Neurog3 expression increased at 24-hr. ChgA and Tph1 increased at 72-hr and peaked at 96-hr (Fig. 2B-D). Consistent with *in vivo* findings, DBZ increased levels of Neurog3, ChgA, and Tph1 in VFKO and NFKO to a similar extent. However, NFKO mGO showed a more time-restricted effect on Neurog3 expression than VFKO (Fig. 2B), while the effect on ChgA and Tph1 was similar (Fig. 2C, D).

**Figure 2.**
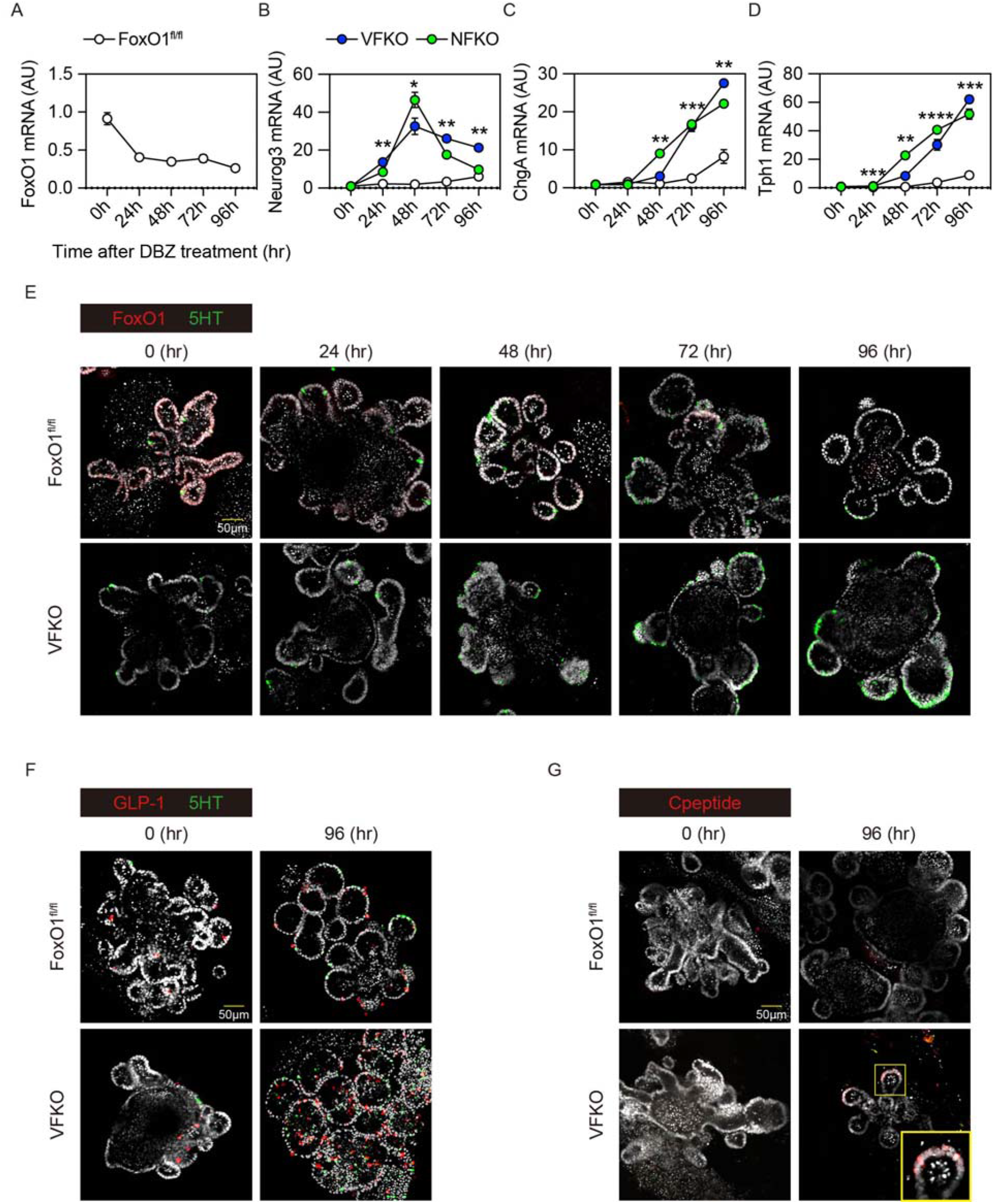
Time course of EEC differentiation after FoxO1 ablation. (A–D) Time course analysis of FoxO1 (A), Ngn3 (B), ChgA (C), and Tph1 (D) expression in mouse organoids after DBZ treatment. All the data were normalized by 18S. White, blue and red filled circles indicate FoxO1^fl/fl^, NFKO, VFKO mice mGO, respectively. (E–G) Immunohistochemistry of FoxO1 and 5HT (E), GLP-1 and 5HT (F), and C-peptide (G) in mGO from FoxO^fl/fl^, or VFKO mice after DBZ treatment. NFKO, Neurog3-Cre (+); FoxO1^fl/fl^: VFKO, Villin-Cre (+); FoxO1^fl/fl^. Scale bar = 50 μm. DAPI counterstains nuclei. Data are presented as means ± SD. *= p < 0.05; **= p < 0.01; ***= p < 0.001; ****= p < 0.0001.

We have shown previously that FoxO1 inhibition increases the number of 5HT cells in human gut organoids (hGO) ^7^. Therefore, we assessed FoxO1- and 5HT-immunoreactive cells during DBZ treatment (Fig. 2E). FoxO1 was expressed in nearly all cells at the beginning of treatment, but levels were suppressed as early as 24-hr especially in crypts, and nearly abolished by 72-hr. In contrast, 5HT cells were increased at 72-hr. In VFKO mGO, 5HT cells increased as early as 48-hr and peaked at 72-hr during DBZ treatment. DBZ treatment increased the number of GLP-1^+^ cell number in mGO isolated from wild type mouse ileum and, again consistent with findings in hGO ^7^, we found a higher number of GLP-1^+^ cells in VFKO mGO at baseline and after DBZ treatment compared to control (Fig. 2F). We next tested the effect of dual inhibition of FoxO1 and Notch on the generation of β-like-cells. We detected C-peptide^+^ cells in DBZ-treated VFKO mGO after 96-hr (Fig. 2G). These results suggest that dual inhibition of FoxO1 and Notch enhanced the number of EEC lineage cells, including GLP-1^+^ cells, and re-directed a subset of cells toward the β-like-cell phenotype.

### Metabolic impact of pan-gut epithelial FoxO1 ablation

We have shown that Neurog3-driven FoxO1 ablation lowers glycemia through effects conveyed by gut β-like-cells in mice rendered diabetic with streptozotocin (STZ) ^6^. To examine the impact of pan-epithelial gut FoxO1 ablation on glucose metabolism, we analyzed VFKO mice by immunohistochemistry and oral (OGTT) or intraperitoneal (ipGTT) glucose tolerance tests. First, we evaluated the impact of pan-intestinal epithelial FoxO1 ablation on gut structure, body weight, and food intake. Unlike a prior study^22^, villin-cre-driven FoxO1 ablation did not affect any of these parameters under our experimental condition (Fig. S3A–D). Consistent with the mGO experiments, we found increased numbers of GLP-1^+^ cells in ileum and no difference in GIP^+^ cells in duodenum compared to controls (Fig3A, B, D, E). We found C-peptide^+^ cells exclusively in VFKO mice (Fig. 3C, F). VFKO mice showed a slight decrease of glucose AUC during OGTT (Fig. S4A and B), but not ipGTT (Fig. S4C and D). Plasma insulin concentrations were significantly higher at 15 and 30 min during OGTT in VFKO compared to controls, presumably due to gut insulin^+^ cells and increased GLP-1^+^ cells (Fig. S4E). To enhance the glucose tolerance phenotype seen in VFKO mice using Notch inhibition, we treated mice by oral dosing of the Notch inhibitor PF-03084014 (hereafter, PF) ^23^ for 4 days. We optimized dosing of PF and found that oral dosing at 150 mg/kg increased EEC number and retained proliferative cells in crypts (Fig S5A–E). Both vehicle and PF treatment did not affect body weight (Fig. 3G and J). PF treatment resulted in a more prominent improvement of glucose tolerance in VFKO compared to vehicle (Fig. 3H, I, K, and L). These data are consistent with a synergistic effect of FoxO1 and Notch dual inhibition to improve glucose tolerance, possibly due to newly generated insulin^+^ and GLP-1^+^ cells.

**Figure 3.**
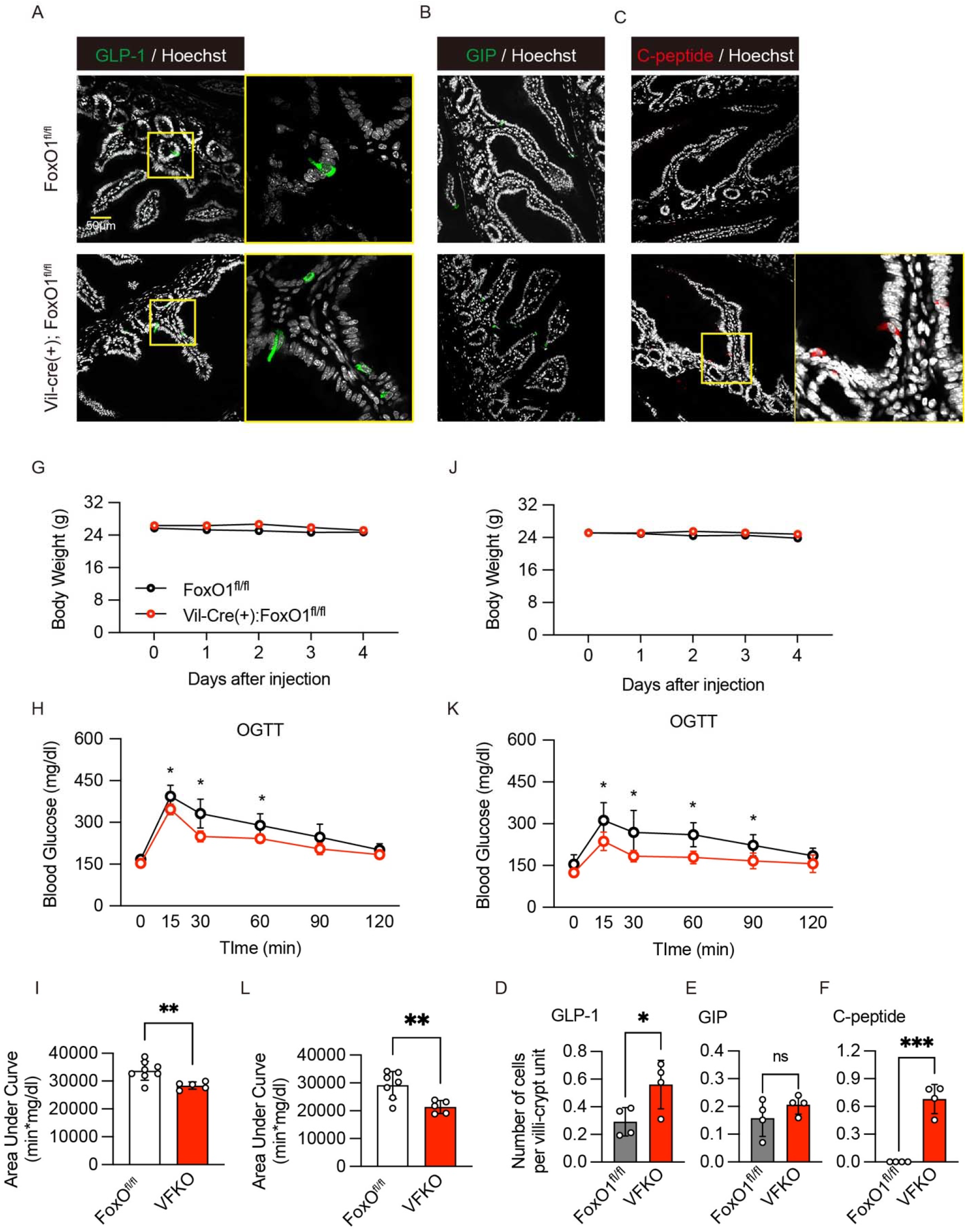
Metabolic effects of gut FoxO1 ablation. (A–B) Immunohistochemistry with GLP-1 (A), GIP (B), and C-peptide antibodies (C) in ileum (A), or duodenum (B and C) of FoxO^fl/fl^ or VFKO mice. The quantified data of (A–C) are shown in (D–F). White and red circles represent FoxO^fl/fl^ and VFKO mice, respectively. VFKO, Vil-Cre (+); FoxO1^fl/fl^: Scale bar = 50 μm. DAPI counterstains nuclei. Data are presented as means ± SD *= p < 0.05; ***= p < 0.001

### Gut FoxO1 ablation reverses hyperglycemia in INS2^Akita/+^

To investigate the glucose-lowering effect associated with induced intestinal β-like cells, we introduced null *FoxO1* alleles in an insulin-deficient diabetic mouse model. Akita mice develop insulin-dependent diabetes by 3- to 4-weeks of age due to a dominant-negative mutation in the *Ins2* gene ^24^. We crossed VFKO mice with INS2^Akita/+^ mice to generate INS2^Akita/+^; Vil-Cre (+): FoxO1^fl/fl^ (Akita-VFKO) and WT littermate controls (INS2^Akita/+^; FoxO1^fl/fl^ (Akita-WT). We monitored glucose levels from 4 to 24 weeks in ad libitum-fed conditions. Both Akita-WT and Akita-VFKO mice showed increased glucose levels starting at 4 weeks, but Akita-VFKO mice showed only mild hyperglycemia and maintained consistently lower glucose level than Akita-WT for the duration of the experiment (Fig. 4A). This effect was confirmed after 4-hr fasting and was not associated with hypoglycemia (Fig. 4B). Plasma insulin levels in ad libitum-fed Akita-VFKO mice were significantly higher compared to Akita-WT starting at 10 weeks of age (Fig. 4C). Moreover, OGTT in 12-week-old Akita-VFKO animals demonstrated significant improvement in glucose tolerance compared to Akita-WT (Fig. 4D). We evaluated whole pancreatic insulin content to rule out a contribution from endogenous pancreatic insulin secretion in FoxO1-ablated animals (Fig. S6A–E). However, the Akita mutation resulted in extreme depletion of pancreatic insulin content regardless of FoxO1 ablation (Fig. S6E). To evaluate whether the improved glucose tolerance was due to increased numbers of GLP-1 cells, we performed OGTT in the presence of the GLP-1 antagonist, exendin-9. Exendin-9 treatment alone increased glucose levels in both groups but had no additional effect in Akita-VFKO mice, indicating that the improvement of OGTT is GLP-1-independent (Fig 4E, F).

**Figure 4.**
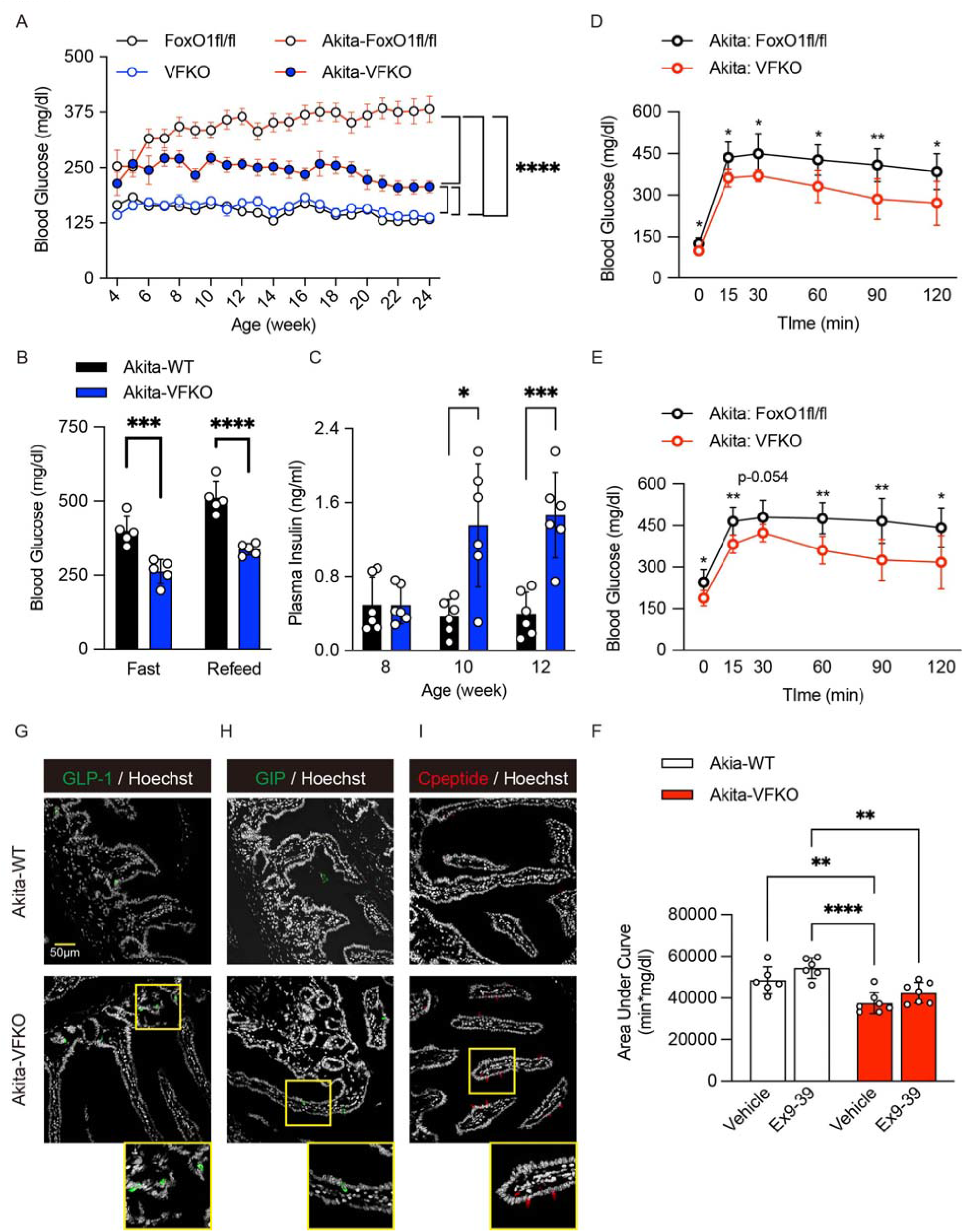
Gut FoxO1 ablation reverses hyperglycemia in INS2^Akita/+^. (A) Blood glucose levels in *INS2^Akita/+^* mice with or without FoxO1^fl/fl^ and VFKO alleles (n = 14 - 20 in each group). (B) Blood glucose levels in *INS2^Akita/+^* and *INS2^Akita/+^*VFKO mice at 16 – 24 weeks ages after 4-hr fast, followed by 1-hr refeeding (n = 5 in each group). (C) Plasma insulin levels in *INS2^Akita/+^* and *INS2^Akita/+^*VFKO mice at 8, 10, and 12 weeks ages (n = 6 in each group). (D-E) Oral glucose (1g/kg glucose infusion) tolerance tests (OGTT) after 16-hr fast of *INS2^Akita/+^* (white circles) and *INS2^Akita/+^*VFKO mice (red circles) following administration of vehicle (D) or GLP-1 antagonist, exendin-9 by i.p. (E) (n = 6 and 7 in each group). (F) Area under the curve (AUC) measurements of OGTT in (D) and (E). (G–I) Immunohistochemistry of GLP-1 (G), GIP (H), and C-peptide (I) in ileum (G), or duodenum (H–I) of *INS2^Akita/+^* and *INS2^Akita/+^*VFKO mice. VFKO, Vil-Cre (+); FoxO1^fl/fl^: Akita-WT, INS2^Akita/+^; FoxO1^fl/fl^: Akita-VFKO, INS2^Akita/+^; Vil-Cre (+): FoxO1^fl/fl^. Data are presented as means ± SD. Scale bar = 50 μm, DAPI counterstains nuclei. *= p < 0.05; **= p < 0.01; ***= p < 0.001 by two-factor ANOVA.

GLP-1-, and C-peptide-immunoreactive cells in duodenum and ileum increased significantly in Akita-VFKO compared to controls (Fig. 4G–I, S6F–H). These data are consistent with the glucose-lowering effect of gut-derived β-like cells observed in STZ-treated Neurog3-FoxO1 mice ^6^.

### Induction of cell conversion by FoxO1 and Notch inhibition

Next, we asked whether a previously described selective FoxO1 inhibitor (compound 10; hereafter, Cpd 10) ^13, 14^ can induce β-like cells in gut organoids and *in vivo*. In primary mGO cultures, Cpd 10 phenocopied the effect of genetic FoxO1 ablation on EEC differentiation, resulting in a time-dependent induction of Neurog3, ChgA, and Tph1 compared to vehicle (Fig. 5A–D). Interestingly, Cpd10 also increased FoxO1 mRNA levels, consistent with the effect of dominant-negative FoxO1 mutants to cause a compensatory activation of endogenous *Foxo1* transcription^7, 25, 26^. These gene expression changes were confirmed by immunohistochemistry with FoxO1 and Tph1 antibodies (Fig. S7A). Cpd10 treatment induced C-peptide-immunoreactive cells but –unlike the genetic ablation– did not induce GLP-1 cells (Fig. S7B, C). To assess the function of these cells, we compared GLP-1 and insulin secretion in mGO from VFKO and NFKO mice or following pharmacological inhibition with Cpd10 in WT mice. Organoids from VFKO mice showed increased basal and glucose-stimulated GLP-1 secretion compared to WT (Fig. 5E). Treatment with DBZ increased basal secretion in control mGO to the levels seen in VFKO and further raised glucose-dependent GLP-1 secretion (Fig. 5E). In contrast, Cpd10 had no effect on GLP-1 secretion in WT mGO, whereas DBZ increased both basal and glucose-stimulated GLP-1 (Fig. 5F). These data are consistent with the immunohistochemistry showing no effect of Cpd10 on the number of GLP-1^+^ cells (Fig. S7B).

**Figure 5.**
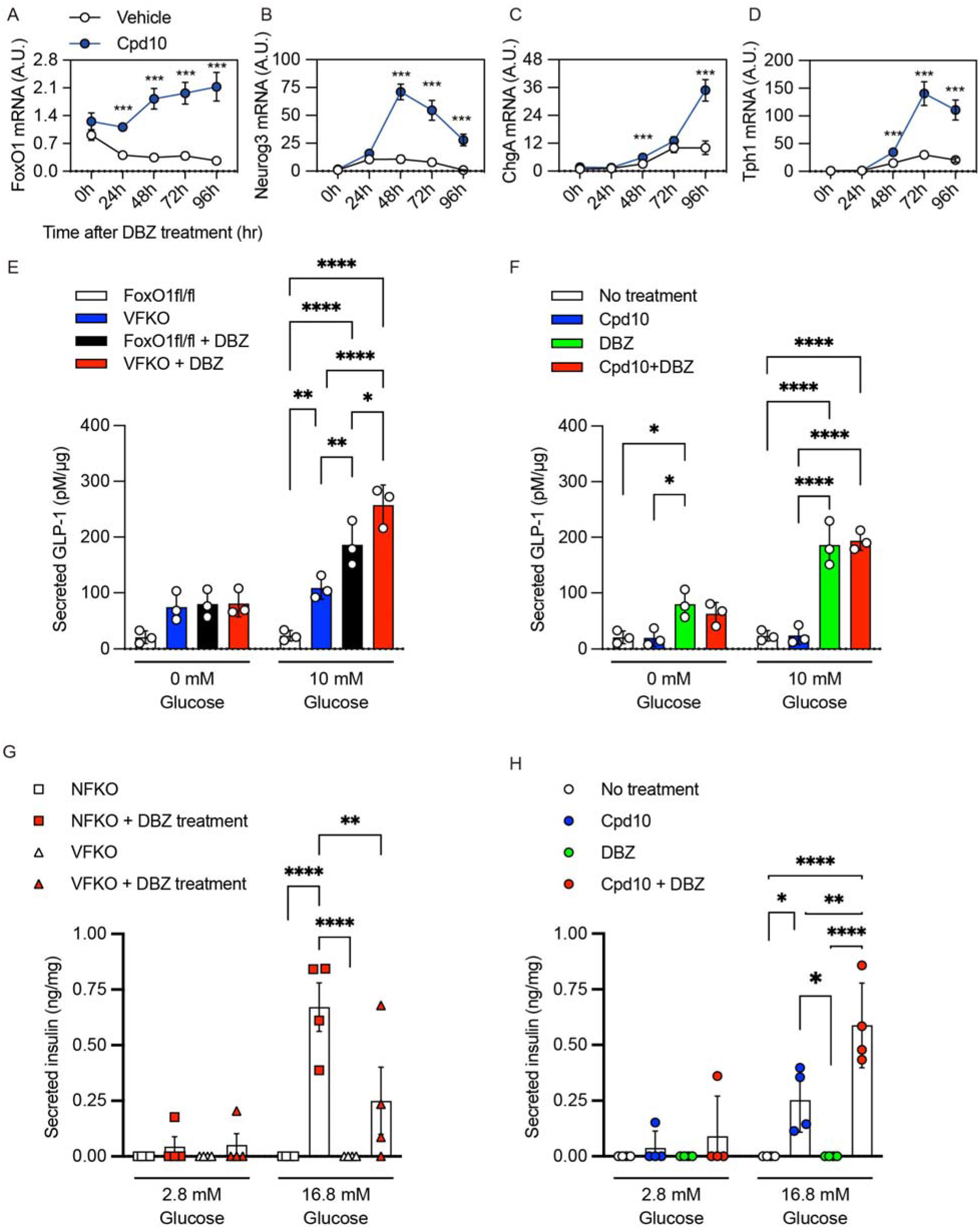
β-like cell conversion by dual FoxO1-Notch inhibition. (A–D) Time course of mGO gene expression following treatment with vehicle (white circles) or Cpd10 (blue circles). (A), Ngn3 (B), ChgA (C), and Tph1 mRNA (D) after DBZ treatment. (E–F) GLP-1 secretion from FoxO1^fl/fl^ or VFKO mice mGO treated with vehicle (white and blue bars) or DBZ (black and red bars) for 96-hr (E). GLP-1 secretion from WT mGO with or without Cpd10 and/or DBZ treatment for 96hr (F). (G–H) Insulin secretion from NFKO or VFKO mGO treated with vehicle (white squares or triangles) or DBZ (red squares or triangles) for 96-hr (G). Insulin secretion from WT mGO with or without Cpd10 and/or DBZ treatment for 96-hr (H). Data are presented as means ± SD. *= p < 0.05; ***= p < 0.001.

mGO derived from NFKO or VFKO mice did not show detectable insulin secretion in response to glucose, likely due to the small number of β-like cells. However, addition of DBZ to the medium resulted in robust glucose-stimulated insulin secretion, ∼ 2.6-fold higher in NFKO than VFKO mGO (Fig. 5G). In WT mGO, Cpd10 alone induced glucose-dependent insulin secretion, and this effect was enhanced by DBZ treatment (Fig. 5H, Fig. S7C). Thus, whereas DBZ has an independent effect to increase GLP-1 secretion ^27^, the augmented insulin secretion effect of DBZ requires either genetic or pharmacologic inhibition of FoxO1.

### FoxO1 and Notch inhibitors lower glucose in INS2^Akita/+^

To evaluate the synergistic effects of FoxO1 and Notch inhibition in the gut *in vivo*, we administered Cpd10 by intraperitoneal injection with or without oral dosing with PF^23^ to INS2^Akita/+^ mice for 5 days. In wild type mice, Cpd10 concentrations in the duodenum peaked within 1-hr of administration compared to plasma and dropped into the low nM range by 24-hr (Fig. S8A). At the employed dose (50mg/kg), Cpd10 did not affect hepatic *G6pc* or *Pck1* expression (Fig. S8B). Combination treatment with 50mg/kg of Cpd10 and 150 mg/kg of PF yielded the highest number of EECs and induced β-cell marker expression (Fig. S9A–B). In addition, the combination treatment significantly increased insulin content in gut epithelial cells from mice (Fig S9C).

Neither single nor combination therapy affected body weight (Fig. 6A). Consistent with the effect of Notch signal inhibition on hepatic glucose production^19^, PF decreased glucose levels (Fig. 6B). Ad lib-fed glucose concentrations decreased by ∼150-170 mg/dl in mice treated with either Cpd10 or PF alone, and a 4-hr-fast did not induce hypoglycemia (Fig. 6C). Combination treatment had no additional benefit on mean glucose concentrations, except in a subset of mice in which glucose levels decreased below 200 mg/dl (Fig. 6B). However, combination treatment had an additive effect to improve OGTT (Fig. 6C-D). Treatment with Cpd10 alone was also associated with increased insulin concentrations (Fig. 6E).

**Figure 6.**
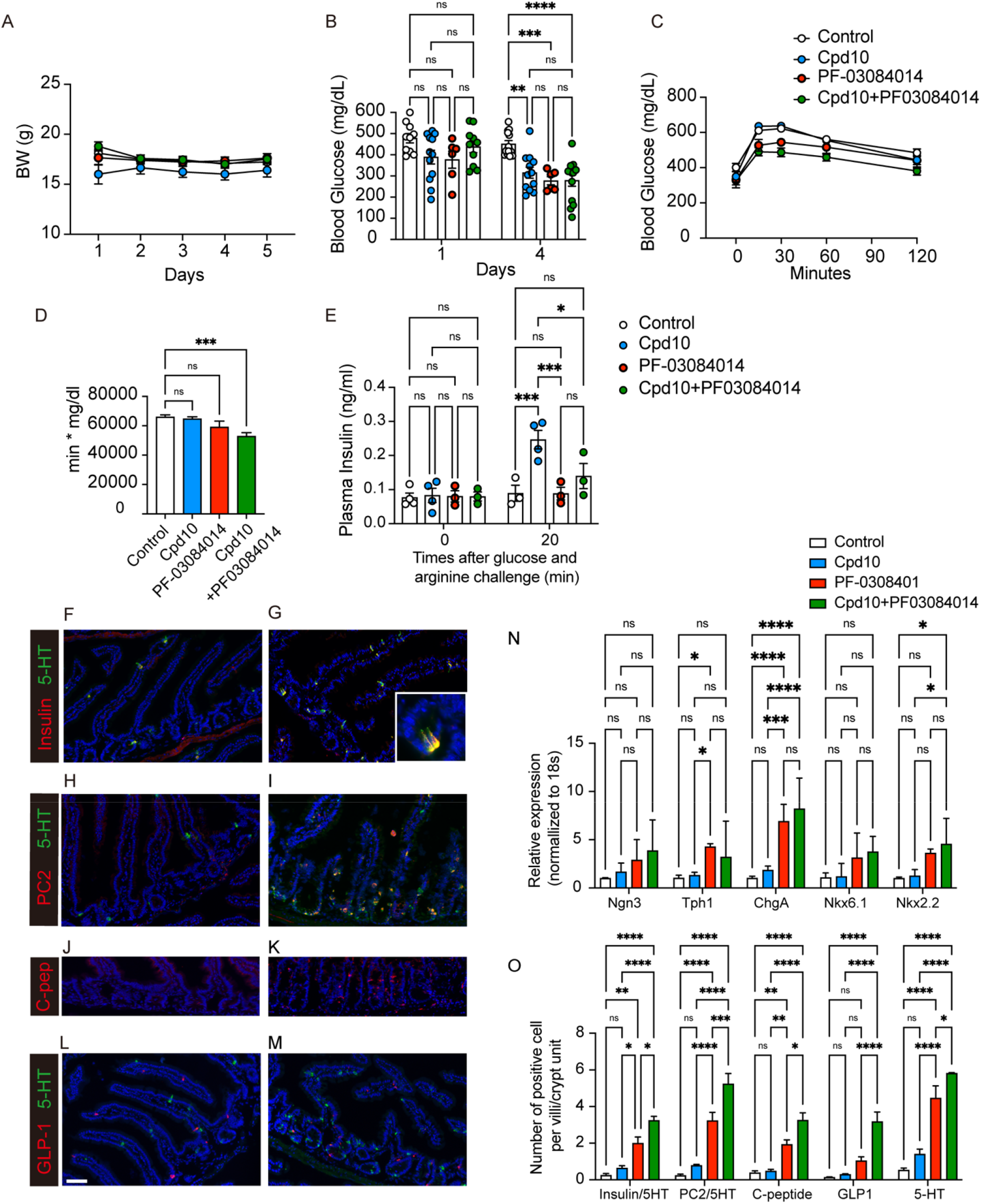
Pharmacological inhibition of Foxo1 and Notch in *AKITA*^*Ins2*/+^ mice. Mice were treated with vehicle, Cpd10 (50 mg/kg/), or PF-03084014 (150 mg/kg) or Cpd10 (50mg/kg) + PF-03084014 (150mg/kg) for 5 days. (A) Body weight measured before morning dosing. (B) glucose levels measured around 5 pm on day 1 and day 4. N= 6-13 per group. (C) Oral glucose tolerance (OGTT) test. 2g/kg of glucose solution was administered orally after a 4-hr fast. (D) Total AUC. (E) Plasma insulin levels at 0 and 20 min after arginine injection. (F-M) Representative intestinal immunohistochemistry after 10-day treatment. (F, G) Insulin (red) and 5-HT (green), (H, I) PC2 (red) and 5-HT (green), (J, K) C-peptide and (L, M) GLP-1 (red) and 5-HT (green) staining of vehicle group (left panel, F, H, J, and L) and Cpd10/PF-03084014-treated group (right panel, G, I, K, M). Scale bar = 50 μm (N) Gene expression in isolated epithelial cells for the markers indicated. N= 4-5 per group. (O) Insulin^+^, PC2^+^, C-peptide^+^, GLP-1^+^, and 5HT^+^ cells quantification in villi and crypts. Data was presented as means ± SEM. *= p < 0.05; **= p < 0.01; ***= p < 0.001; ****= p < 0.0001; NS: not significant. Two-way ANOVA was performed for (B), (E), (N), and (O). One-way ANOVA was performed for (D).

Combination treatment had an additive effect on Neurog3, ChgA, Nkx6.1, and Nkx2.2 mRNA in duodenum, whereas PF alone was sufficient to achieve maximal induction of Tph1 mRNA (Fig. 6N).

Immunohistochemistry demonstrated a synergistic effect of combination treatment to generate up to ∼3.2 C-peptide^+^ and ∼3.2 insulin^+^ cells per villus-crypt unit (Fig. 6F–K, O, S10A–F, S11A–E). PF alone increased the number of GLP-1^+^ cell numbers (Fig. 6L–M, S10G–H).

## DISCUSSION

The present study demonstrates that pharmacological inhibition of FoxO1 can induce β-like cells in the gut similar to genetic FoxO1 knockout. This remarkable property of small molecule FoxO1 inhibitors has not been previously reported. This finding expands on work showing that Cpd10 can reduce hepatic glucose production from pyruvate *in vivo* ^13^. Successfully applied to humans with diabetes, this strategy would provide an alternative to insulin injections, circumventing the risk of hypoglycemia and eliminating the need for frequent glucose monitoring, thus significantly reducing the burden of care and potentially improving glycemic control. Whether gut-derived β-like cells would be targeted by autoimmunity remains a concern. However, based on the distinct immunity of the gut mucosa compared to the pancreas ^9^, rapid cell turnover, and scattered distribution of β-like cells, it can be argued that they can escape functionally consequential autoimmune engagement ^28^.

Since the original report on the role of FoxO1 in the gut ^6^, the impact of FoxO1 ablation on trans-differentiation of entero-endocrine cells (EECs) has been confirmed in human gut organoids ^7^. Experimental co-expression of Neurog3, Pdx1, and MafA can generate β-like cells in the gut ^5^. FoxO1 inhibition recapitulates these transcriptional effects ^29, 30^. FoxO1 activation can suppress Neurog3 ^31^, possibly via O-GlcNAc transferase resulting in fewer GLP-1 cells ^32^.

### Mechanism of cell conversion and combination with Notch

To define the stage in EEC differentiation affected by FoxO1 ablation, we show that FoxO1 inactivation in epithelial progenitors (VFKO) phenocopies inactivation in the EEC progenitor (NFKO), consistent with a primary role of FoxO1 in EEC progenitors to suppress the β-like-cell phenotype. We also identified two developmental stages in the EEC progenitors by dual tracing with GFP and RFP, and tentatively mapped the role of FoxO1 inhibition to the late progenitor/early EEC stage. This narrow window of FoxO1 action may explain why the yield of insulin-immunoreactive cells is comparatively lower in FoxO1 knockout models than in the triple transcription factor gain-of-function transgenics ^33, 34^.

Leveraging an extensive prior body of work regarding FoxO1 and Notch to regulate endocrine cell lineage ^15, 35, 36^, we successfully tested a combination regimen that increased the number of β-like cells. Notch inhibition promoted expansion of the EEC lineage, truncating the stage between Lgr5^+^ and Neurog3^+^. By expanding the pool of available late progenitors, we saw an increased generation of β-like cells due to FoxO1 inhibition. This interplay was confirmed over a 96hr treatment of primary organoids. We also detected suggestive evidence of an additive glucose-lowering effect in a subset of Akita mice treated with the combination of Cpd10 and PF. The increased supply of insulin from gut-derived β-like cells induced by Cpd10 alone reduced hyperglycemia in Akita mice. Combination treatment enhanced this effect. Permutations of the dosing regimen (e.g., route of delivery, pulse-chasing, timed delivery, alternate day) will be needed to explore the full glucose-lowering potential of the combination.

There are differences between genetic ablation and chemical inhibition of FoxO1. Chemical inhibition was applied to mGO at various stages, and only cells at the Ngn3-expressing stage could be differentiated into β-like cells. In addition, Villin-driven FoxO1 ablation increased both GLP-1 and β-like cells, while Cpd10 increased insulin, but not GLP-1 secretion. These differences may reflect selective inhibition of FoxO1 targets by Cpd10, as shown in prior work in which this compound demonstrated selective inhibition of expression of G6pc but not Gck in liver^14^. A similar situation may contribute to the different effects on generation of GLP-1-expressing cells. Cpd10 may specifically inhibit FoxO1 targets involved in cellular lineage specification. Development of additional FoxO1 inhibitors may enable differential targeting of the EEC lineage-promoting and β-like cell-promoting functions.

### Risks and limitations

We are aware of the risks and limitations of the approaches used here. While there is growing interest in FoxO proteins as drug targets ^37^, the gamut of their functions makes prediction of potential adverse effects daunting. Two areas of concern are carcinogenesis ^38^ and autoimmunity ^39^. Inducible post-natal deletion of FoxO1, 3, and 4 resulted in thymic lymphomas and hemangiomas ^38^. However, this phenotype requires inactivation of all three isoforms, while our inhibitors are restricted to FoxO1, a counter-screen against FoxO3 being part of the developmental pipeline ^13, 14^. Furthermore, evidence linking FoxO1 loss-of-function to cancer is sparse and contradictory, while somatic FoxO1 *<u>gain</u>*-of-function is associated with forms of non-Hodgkin lymphoma^40, 41^ suggesting that FoxO1 inhibitors could indeed be beneficial in some cancers. Furthermore, FoxO1 is not expressed in gut epithelial stem cells, and inhibitors can be designed to provide minimal systemic exposure, further lowering the risk. We did not see gut neoplasia in VFKO mice of up to 24 weeks of age. Integrating these insights, gut-restricted FoxO1 inhibition is unlikely to incur carcinogenic risks.

Another potential adverse effect of FoxO1 inhibition is autoimmunity, which would be problematic in type 1 diabetes ^39^. Evidence of FoxO1 deletion causing autoimmunity comes exclusively from genetic knockouts in highly selective cell types, such as Treg cells ^39^. Unlike the widely agreed metabolic functions of FoxO1, the immune functions of FoxO1 are highly context-dependent, including evidence of protecting against autoimmunity in gut and airway ^42, 43^. Given the multiplicity of roles assigned to FoxO1, and the apparent safety of gut inhibition, future treatments should aim for a gut-restricted compound with minimal systemic exposure.

Notch inhibitors have been tested in Alzheimer’s disease and cancer, generally with modest efficacy but excellent safety profiles; modest gastrointestinal upset due presumably to goblet cell hyperplasia has been reported ^44^. Whether the combination with FoxO1 inhibitors will increase such effects remains to be seen. However, given the turnover rate of the gut epithelium, any gastrointestinal adverse effects are expected to be rapidly reversible after cessation of treatment, and this would be formally assessed in preclinical safety studies in due course. It should also be pointed out that a clinical development path for a type 1 diabetes indication is only marginally affected by the need to test compounds alone and in combination, since clinical efficacy can be determined swiftly with a limited number of subjects ^45^. In the present *in vivo* studies, we used an early generation Notch inhibitor with anti-hepatocellular carcinoma effects in mice ^23^. The antihyperglycemic effect of dual inhibition was evident after 5 days of treatment.

The present study paves the way to develop an oral treatment to replace insulin via gut-derived insulin-secreting cells for type 1 diabetes patients. While the present compound combination provides proof-of-principle, further combinations of FoxO1 and Notch inhibitors should be explored to maximize glucose-lowering effects and de-risk toxicity and safety. Success would result in a paradigm-shifting strategy to replace insulin administration in type 1 diabetes.

## Materials and Methods

### Animals

Mice were housed in a climate-controlled room on a 12h light/dark cycle with lights on at 07:00 and off at 19:00, and were fed standard chow (PicoLab rodent diet20, 5053; PurinaMills). Male mice of C57BL/6J background aged 8-12 weeks were used. To generate enteroendocrine- and epithelial gut-specific FoxO1 knockouts, we crossed FoxO1lox/lox and Tg (Neurog3-cre)C1Able/J (Ngn3-Cre)^20^ or B6.Cg-Tg(Vil1-cre)1000Gum/J (Vil-Cre) ^21^ transgenic mice. For lineage tracing and cell sorting, we crossed Gt(ROSA)26Sortm4(ACTB-tdTomato,-EGFP)Luo/J (mTmG)^46^ with Ngn3-Cre mice. For diabetic mice model, we used C57BL/6-Ins2Akita/J (Akita) ^24^. To assess the time course of blood glucose levels with/without FoxO1 knockouts in Akita mice, we measured blood glucose levels between 09:00 to 10:00 in ad libitum condition. Blood glucose was measured from tail clip capillary blood using (CONTOUR®NEXT ONE, Ascensia, USA). In the Enteroendocrine induction model, we injected Gamma-secretase inhibitor (DBZ, 25 mpk dissolved in 0.5% Methocel / 0.1% Tween80) (Chiralix, Netherlands) at 10:00 for two consecutive days. Tissues were harvested 96-hr after the first injection and analyzed by immunohistochemistry. All animal experiments were in accordance with NIH guidelines, approved and overseen by the Columbia University Institutional Animal Care and Use Committee.

### Metabolic testing

We used male mice aged 8–12 weeks. We performed intraperitoneal or oral glucose tolerance tests after a 4h- (10 a.m. to 2 p.m.) or 16h- (6 p.m. to 10 a.m.) fast administering 1 to 2 g/kg glucose infusion by i.p. or p.o. Blood glucose and insulin measurements were made from tail vein blood. Insulin was measured with a mouse-specific ELISA kit (Mercodia, USA). For glucose tolerance test with pharmacological GLP-1 blockade, 10 μg/100 μl per mouse of Exendin (9-39) (Bachem, CA) or vehicle (saline) were injected by i.p. 20 minutes before glucose measurements.

### Drug treatment

6- to 7-week-old male Akita or C57BL/6J mice were acclimated for two weeks and randomized according to body weight and ad libitum glucose levels. Cpd10 was formulated in 5% Kolliphor® HS 15 /water for i.p injection or N,N-Dimethylacetamide:Kolliphor® HS 15: water=5:10:85 (v/v/v) solution, pH4∼5 for oral gavage injection. PF-03084014 was formulated in N,N-Dimethylacetamide:Kolliphor® HS 15: water=5:10:85 (v/v/v) solution, pH4∼5. Mice were dosed at 10 ml/kg twice with Cpd10 by i.p., and once with PF-03084014 daily for 5-15 days. Body weight was monitored daily in the morning before the first dose, and ad libitum blood glucose levels were monitored every 2-3 days using Contour Next glucometer or StatStrip Xpress (Novamedical) from tail vein. Oral glucose tolerance test (OGTT) was performed 5 days after treatment, by gavaging 2 g/kg (10 ml/kg) of D-glucose (Sigma) dissolved in the distilled water after 4-hr fasting from 8:30 am to 12:30 pm. Tail blood glucose was measured at 0, 15, 30, 60 and 120 min. Mice were dosed 2-hr prior to injecting glucose.

### Chemicals

Ketamine was from KetaSet® and Xylazine from AnaSed®; Ketamine was from KetaSet® and Xylazine from AnaSed®; DBZ from Chiralix, PF-03084014 (Nirogacestat) from SelleckChem; Cpd10 from ForkheadBioTherapeutics, and Kolliphor® HS 15 from MilliporeSigma; N,N-Dimethylacetamide from MilliporeSigma. Advanced DMEM/F12, HBSS, EDTA, HEPES were from Invitrogen; Bovine Serum Albumin (BSA) from Fisher Scientific. 16% paraformaldehyde (PFA) was from Electron Microscopy Sciences and was diluted in sterile phosphate buffer solution to 4% final concentration.

### Drug exposure test

8-week-old C57BL6/J mice were used for drug exposure experiments. Mice were fasted for 2-hr, and 10mg/kg Cpd10 was administered by gavage. Blood was collected and small intestine harvested 1-, 4-, 8-, and 24-hr after dosing. The samples were treated with acetonitrile/methanol (1:1 v/v), centrifuged and analyzed by LC/MS (pump: NEXERA LC-30AD; controller: Shimadzu CBM-20A; autosampler: PAL System; mass spectrometer: ABSciex API4000; column: Advantage ECHELON C18 4 µm 20 × 2.1mm (SpriteAE1822)) using a gradient (phase A: Water/1M NH4HCO3, 2000:10, v/v; phase B: MeOH/1M NH4HCO3, 2000:10, v/v). Results were confirmed by biological duplicates at each time points.

### RNA isolation and quantitative PCR analysis

We lysed whole gut, FAC-sorted cells (5-10 x 10^3^), or mGO in 1 mL TRIzol, purified RNA using RNeasy Mini Kit (Qiagen, Germantown, MD), reverse-transcribed it with qScript cDNA Synthesis Kit (QuantaBio, Beverly, MA), and performed PCR with GoTaq® qPCR Master Mix (Promega, Madison, WI). PCR primers are listed in Table S1. Gene expression levels were normalized to 18S using the 2-DDCt method ^47^ and are presented as relative transcript levels.

### Immunohistochemistry and histomorphometry

We anesthetized 8- to 12-week-old mice and harvested small and large intestine and rinsed them in PBS. Swiss rolls were prepared by rolling up the intestines around a pick stick from the pyloric sphincter toward cecum for small intestine, and from rectum to cecum for colon. Preparations were collected, fixed in 4% paraformaldehyde for 2-hr, dehydrated in 30% sucrose overnight at 4°C, embedded in OCT (Sakura, Torrance, CA), frozen to −80°C, and cut into 7-µm sections. The primary and secondary antibodies used are listed in Table S2. Insulin and FoxO1 staining for gut tissues were visualized by the Avidin/Biotin Blocking Kit (Vector Laboratories, CA) and biotinylated secondary antibody (BD, NJ) and fluorochrome-conjugated streptavidin (Vector Laboratories, CA). For Cpeptide staining, we treated samples for antigen retrieval by boiling in citrated buffer at 70° for 20 minutes (Histo VT One, Nacalai USA, CA).

Images were obtained with a confocal laser-scanning microscope (LSM 710, Carl Zeiss, NY) using Zen software. Quantification was performed using ImageJ software [National Institutes of Health]. The number of stained cells in the intestinal epithelium was counted manually using the Adobe Photoshop 2021 software. The whole section of swiss roll was scanned under 20x magnification using the Keyence BZ-X800 microscope. Images were stitched using the Keyence Analyzer software. 200-300 villi-crypt unit or crypts were marked and counted on the image of nuclei staining, followed by merging the image of insulin or C-peptide staining and counting the positive signals only in the marked villi-crypt unit or crypts.

### Fluorescence-Activated Cell Sorting

The tissue handling from murine intestinal epithelial cells isolation to fluorescence-activated cell-sorted analysis has been described ^48^ and was used in these experiments with slight changes. A segment of the proximal intestine representing a 5-cm region from 2 to 7 cm distal to the pyloric sphincter was used for cell isolation. The intestinal segment was washed, longitudinally cut open, and lightly rinsed to remove residual luminal contents. Tissue was incubated in PBS containing 30 mmol/l EDTA and 1.5 mmol/l dithiothreitol on ice for 20 min. Intestinal tissue was transferred to a 15-ml conical tube containing 5 ml of 30 mmol/l EDTA made in PBS, incubated at 37°C for 8 min, then shaken by hand along the tube’s long axis. After shaking, the remnant muscle layer was removed and the dissociated cells were pelleted at 1,000 g, resuspended in PBS supplemented with 10% FBS, and centrifuged again. Cells were incubated in 10 ml of modified HBSS containing 0.3 U/ml of dispase (Stem Cells Inc., CA) at 37°C for 10min with intermittent shaking every 2 min for 15s to dissociate epithelial sheets to single cells. After 10min, 1 ml of FBS and 1,000 U DNase I (Sigma) were added to cell suspension, which was sequentially passed through 70-μm then 40-μm filters (Fisher Scientific, NH). Cells were pelleted, then resuspended in 10 ml of HBSS to inactivate the dispase. Cells were centrifuged and resuspended at a concentration of ∼2 × 10^7^ cells/ml in FACS buffer [HBSS, EDTA (1mM), HEPES (25mM), 1% BSA, Penicillin-Streptomycin]. SYTOX™ Red was used for the detection of dead cells (Invitrogen, MA). Cells were sorted using a BD Influx with a blue (488-nm) laser and 530/40 band pass filter, green (532-nm) laser and 580/30 band pass filter, red (640-nm) laser and 670/30 band pass filter to detect endogneous Green, Red proteins, and Sytox Red. RNA was isolated from 5,000 – 10,000 sorted cells, following to perform quantitative PCR analysis. To analyze the population of enteroendocrine cells, the isolated cells were first stained with live/dead cell staining kit (L34964), washed with 1x PBS and fixed in BD CytofixTM fixation buffer (BD Bioscience 554655) for 20 min. Cells were then permeabilized in permeabilization buffer (0.2% saponin, 0.5% BSA and 2mM EDTA in PBS) for 10 min. The cells were incubated with the indicated primary antibodies diluted in the permeabilization buffer for 1hr at 4°C, washed 3 times with permeabilization buffer, followed by the secondary antibodies diluted in permeabilization buffer for 30 min at 4 °C. After 3 times of washes, 1 × 10^5^ cells were analyzed using BD LSR FortessaTM Cell Analyzer.

### Pancreatic and gut tissue insulin content

To assess insulin content, the whole pancreas or gut tissue was removed rapidly and extracted in ice-cold acid ethanol overnight at −20°C. Then, the samples are homogenized and incubated overnight at −20°C. The supernatant added with the equivalent volume of ice-cold 1M Tris was used for the analysis. The insulin contents in the samples were determined by mouse insulin ELISA (Mercodia).

### Gut Organoid cultures

Mouse duodenum or ileum crypts were isolated. Duodenal samples included a 5-cm segment of intestine 2 to 7 cm distal to the pyloric sphincter, ileum samples 2 to 7 cm proximal to the cecum, as described previously ^49^. We seeded crypts into Matrigel (Corning, NY) to form mouse gut organoid (mGO) and cultured them for 48-hr prior to experiments. Basic medium was composed of Advanced DMEM/F12 containing 2 mM Glutamax, 10 mM HEPES, 100 U/ml penicillin/streptomycin, primocin (100μg/ml) (Invivogen, CA). Complete growth medium was prepared by adding murine EGF (50ng/ml) (Gibco, TX), R-spondin-1 (conditioned medium 1ml/50ml) (gift by Akifumi Ootani, Stanford University, CA), Noggin (100ng/mL) (PeproTech, Inc, NJ), N-acetyl-L-cysteine (1mM) (Millipore Sigma, MA), B27 (1×)(Gibco, TX) to basic medium. The medium was replaced every 48-hr. All the experiments were done within 7 days after plating and done within 5 passages. For passage, every 7 days Matrigel was lysed with Cultrex™ Organoid Harvesting Solution (R&D Systems™, MN), and isolated mGO, mechanically dissociated by pipetting, pelleted and re-plated in fresh Matrigel in 24-well plates at a 1:5 splitting ratio. Three hours treatment with DBZ (5μM) (Chiralix, Netherlands) was used to promote EEC differentiation, followed by replacing the medium with complete growth medium. Cpd10 (1μM) (Forkhead BioTherapeutics, NY) was used for single or dual treatment with DBZ, and Cpd10 was replaced every 48-hr.

### GLP-1 secretion

Mouse ileum organoids were used for GLP-1 secretion assays. Hanks Buffered Salt Solution supplemented with 10mM HEPES, 0.1% fatty acid-free bovine serum albumin was used as basic medium. mGO from 24-well plates were collected in 1.7ml tubes and incubated for 2-hr in basic medium, washed and incubated in 50μl of basic medium with DPP4 inhibitor (50μM) for 1h. After collecting the supernatant, 10mM glucose solution in basic medium with DPP4 inhibitor was added to the tube, and incubated for 1h, after which we collected the supernatant. GLP-1 concentrations were determined by mouse GLP-1 ELISA Kit (Crystal Chem). mGOs were lysed in CelLytic™ M (Sigma-Aldrich) for protein extraction, and protein content was normalized by total protein content quantified with BCA assay (Thermo scientific).

### Insulin secretion

We collected mGO from 24-well plates, and placed them in ice-cold KRBH buffer (119 mM NaCl, 2.5 mM CaCl2, 115 1.19 mM KH2PO4, 1.19 mM Mg2SO4, 10 mM HEPES pH 7.4, 2% BSA, and 11.6 or 2.8 mM glucose) and incubated at 37°C for 1-hr, followed by incubation in 2.8 mM glucose for 1-hr at 37°C. After collecting the supernatant, 16.8mM glucose was added, and mGO incubated for 1-hr, followed by collection of the supernatant. Insulin was determined by mouse insulin ELISA (Mercodia). mGOs were lysed in CelLytic™ M (Sigma-Aldrich) for protein extraction, and protein content normalized as above.

### Statistical Analysis

Values are presented as means ± standard deviation, and analyzed using Prism 8.2.1 (GraphPad Software, Inc.). We used unpaired Student’s *t*-test for normally distributed variables for comparisons between two groups, one-way ANOVA followed by Bonferroni post-hoc test for multiple comparisons, and Pearson’s correlation coefficient to investigate the relationship between two variables. We used a threshold of *p_alpha (2-tailed)_* < 0.05 to declare statistical significance.

## List of Supplementary Materials

Table S1. List of primers for qPCR

Table S2. List of antibodies for immunohistochemistry

Figure S1. Comparisons of progenitor marker expression among subgroups of sorted EECs by RFP and GFP related to Figure 1

Figure S2. Induction of EECs by DBZ related to Figure 1

Figure S3. Gut structure, weight, and food intake in Vil-FoxO1 knockout and wild-type control mice related to Figure 3

Figure S4. ipGTT and OGTT in Vil-FoxO1 knockout and wild-type control mice related to Figure 3

Figure S5. Optimization of in vivo PF-03084014 treatment related to Figure 3

Figure S6. Comparison of pancreatic insulin content and quantification of GLP-1+, GIP+, and C-peptide+ cells in INS2Akita/+ mice with or without pan-intestinal FoxO1 ablation related to Figure 4

Figure S7. Effect of Cpd10 on FoxO1, 5HT, GLP-1, and C-peptide expression related to Figure 5

Figure S8. Pharmacological effect of Cpd10 in vivo related to Figure 6

Figure S9. Cpd10 and PF-03084014 dual treatment increases enteroendocrine cells related to Figure 6

Figure S10. Representative intestinal Immunohistochemistry after 10-day single treatment with Cpd10 or PF-03084014 related to Figure 6

Figure S11. Images of β-cell specific markers after 10-day treatment related to Figure 6

## Author Contributions

Conceptualization: T.K, D.A

Methodology: T.K, Y.K.L, W.D, S.B

Investigation: T.K, Y.K.L, N.S, B.D, W.M.M, W.D., H.W.

Visualization: T.K, Y.K.L

Funding acquisition: H.L, S.B, D.A

Project administration: T.K, D.A

Supervision: D.A

Writing – original draft: T.K, D.A

Writing – review & editing: T.K, R.L, S.B, D.A, H.L, Y.K.L

## Acknowledgments

We thank members of the Accili laboratory, Dr. Utpal B. Pajvani for insightful discussions of the data, and Mr. Thomas Kolar and Ms. Ana Maria Flete (Columbia University), and Carmen Lam, Kasia Dover, Xiaoming Xu for exceptional technical support.

