## Supplemental file for "Chemical induction of gut β-like-cells by combined FoxO1/Notch inhibition as a glucose-lowering treatment for diabetes"

Running title: FoxO1 inhibitors in murine diabetes

^†^ Corresponding Author: Takumi Kitamoto, MD, PhD

Department of Medicine, Vagelos College of Physicians and Surgeons, Columbia

University, New York, NY

* These authors were equally contributed

Reprint requests: Domenico Accili

INFORMATION

This appendix has been provided by the authors to give extended information about the present study.

Table S1. List of primers for qPCR

Table S2. List of antibodies for immunohistochemistry

Figure S1. Comparisons of progenitor marker expression among subgroups of sorted EECs by RFP and GFP related to Figure 1

Figure S2. Induction of EECs by DBZ related to Figure 1

Figure S3. Gut structure, weight, and food intake in Vil-FoxO1 knockout and wild-type control mice related to Figure 3

Figure S4. ipGTT and OGTT in Vil-FoxO1 knockout and wild-type control mice related to Figure 3

Figure S5. Optimization of in vivo PF-03084014 treatment related to Figure 3

Figure S6. Comparison of pancreatic insulin content and quantification of GLP-1+, GIP+, and C-peptide+ cells in INS2Akita/+ mice with or without pan-intestinal FoxO1 ablation related to Figure 4

Figure S7. Effect of Cpd10 on FoxO1, 5HT, GLP-1, and C-peptide expression related to Figure 5

Figure S8. Pharmacological effect of Cpd10 in vivo related to Figure 6

Figure S9. Cpd10 and PF-03084014 dual treatment increases enteroendocrine cells related to Figure 6

Figure S10. Representative intestinal Immunohistochemistry after 10-day single treatment with Cpd10 or PF-03084014 related to Figure 6

Figure S11. Images of β-cell specific markers after 10-day treatment related to Figure 6

Table S1. List of primers for qPCR

| Gene symbol | | Primer Sequence |
| --- | --- | --- |
| *18S* | Forward | AAACGGCTACCACATCCAAG |
|  | Reverse | CCTCCAATGGATCCTCGTTA |
| *Foxo1* | Forward | TCC AGT TCC TTC ATT CTG CAC T |
|  | Reverse | GCGTGCCCTACTTCAAGGATAA |
| *Ngn3* | Forward | TGGCCCATAGATGATGTTCG |
|  | Reverse | AGAAGGCAGATCACCTTCGTG |
| *ChgA* | Forward | CAGCTCGTCCACTCTTTCCG |
|  | Reverse | CCTCTCGTCTCCTTGGAGGG |
| *Tph1* | Forward | TTCTGACCTGGACTTCTGCG |
|  | Reverse | GGGGTCCCCATGTTTGTAGT |
| *Gcg* | Forward | GCTGATTCAAACCAAGATCACTG |
|  | Reverse | ATCCCAAGTGACTGGCACGAG |
| *Ins1* | Forward | TTAATGGGCCAAACAGCAAAGTC |
|  | Reverse | TGACCTGCTTGCTGATGGTCTC |
| *Ins2* | Forward | CTGGTGGAGGCTCTCTACCTG |
|  | Reverse | CAAGGTCTGAAGGTCACCTGC |
| *Nkx6.1* | Forward | CAGGACCAAGTGGAGAAAGAAG |
|  | Reverse | GGGTCCAGAGGTTTGTTGTAAT |
| *Nkx2.2* | Forward | CCA GAA CCA TCG CTA CAA GAT |
|  | Reverse | CTTATCCAATCGCTCCACCTT |

| Name | Vendor | Cat.No | Primary conc | Secondary conc |
| --- | --- | --- | --- | --- |
| FoxO1 (C29H4) | Cell signaling | #2880 | 1:300 | 1:500 |
| GLP-1 | Abcam | ab13329 | 1:500 | 1:500 |
| GIP | ABBIOTEC | 250668 | 1:500 | 1:500 |
| 5-HT | Abcam | ab66047 | 1:1000 | 1:500 |
| Somatostatin | Dako | A0566 | 1:500 | 1:500 |
| GFP | Abcam | Ab13970 | 1:500 | 1:500 |
| RFP | MBL | PM005 | 1:500 | 1:500 |
| Ki67 | Abcam | ab15580 | 1:500 | 1:500 |
| Insulin | Mercodia | N/A | 1:200 | 1:500 |
| C-peptide | Thermo | PA5-85595 | 1:2000 | 1:500 |
| PC2 | Thermo | PA1-058 | 1:500 | 1:500 |
| Nkx6.1 | R&D | AF5857-SP | 1:50 | 1:500 |
| Donkey anti-goat-Alexa488 | Invitrogen | A11055 |  |  |
| Donkey anti-rabbit-Alexa555 | Invitrogen | A31572 |  |  |
| Donkey anti-rat-Alexa488 | Invitrogen | A21208 |  |  |
| Goat anti-Rabbit-Alexa488 | Invitrogen | A11008 |  |  |
| Goat anti-Mouse-Alexa488 | Invitrogen | A11029 |  |  |
| Goat anti-Mouse-Alexa555 | Invitrogen | A21424 |  |  |
| Goat anti-Chicken-Alexa488 | Invitrogen | A11039 |  |  |
| Biotin rat anti mouse igG1 | BD | 553441 |  |  |

Table S2. List of antibodies for immunohistochemistry

SUPPLEMENTAL FIGURES

Figure S1. Comparisons of progenitor marker expression among subgroups of sorted EECs by RFP and GFP

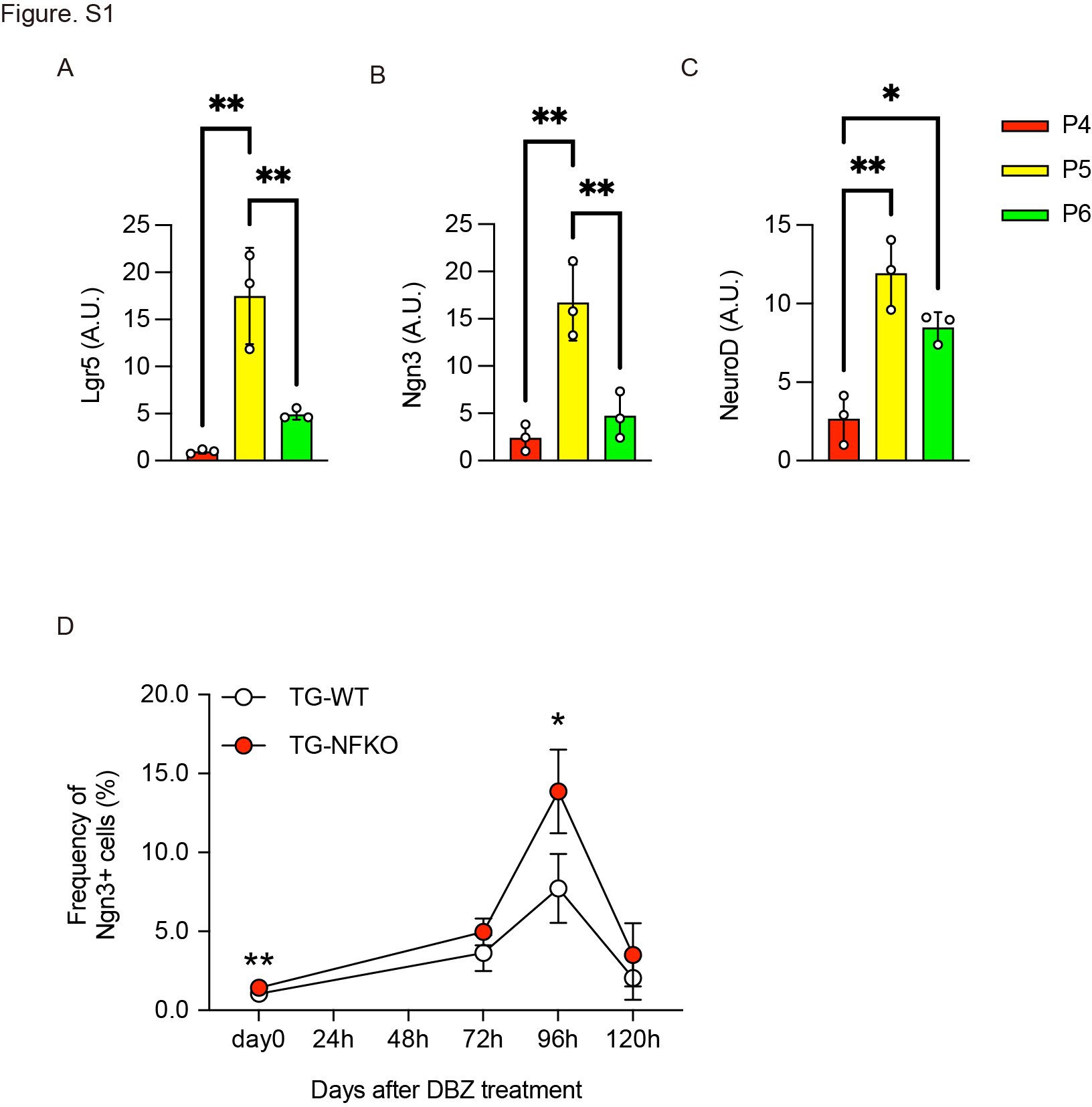

Fig. S1. (A–C) qPCR of Lgr5 (A), Ngn3 (B), and NeuroD (C) using mRNA from sorted P4, P5, and P6 cells indicated in Fig. 1C. N=3 for each group. (D) Time course of total GFP+ cells, indicating Ngn3^+^ EECs after DBZ treatment. The data are calculated from the same samples shown in Fig. 1G–J.

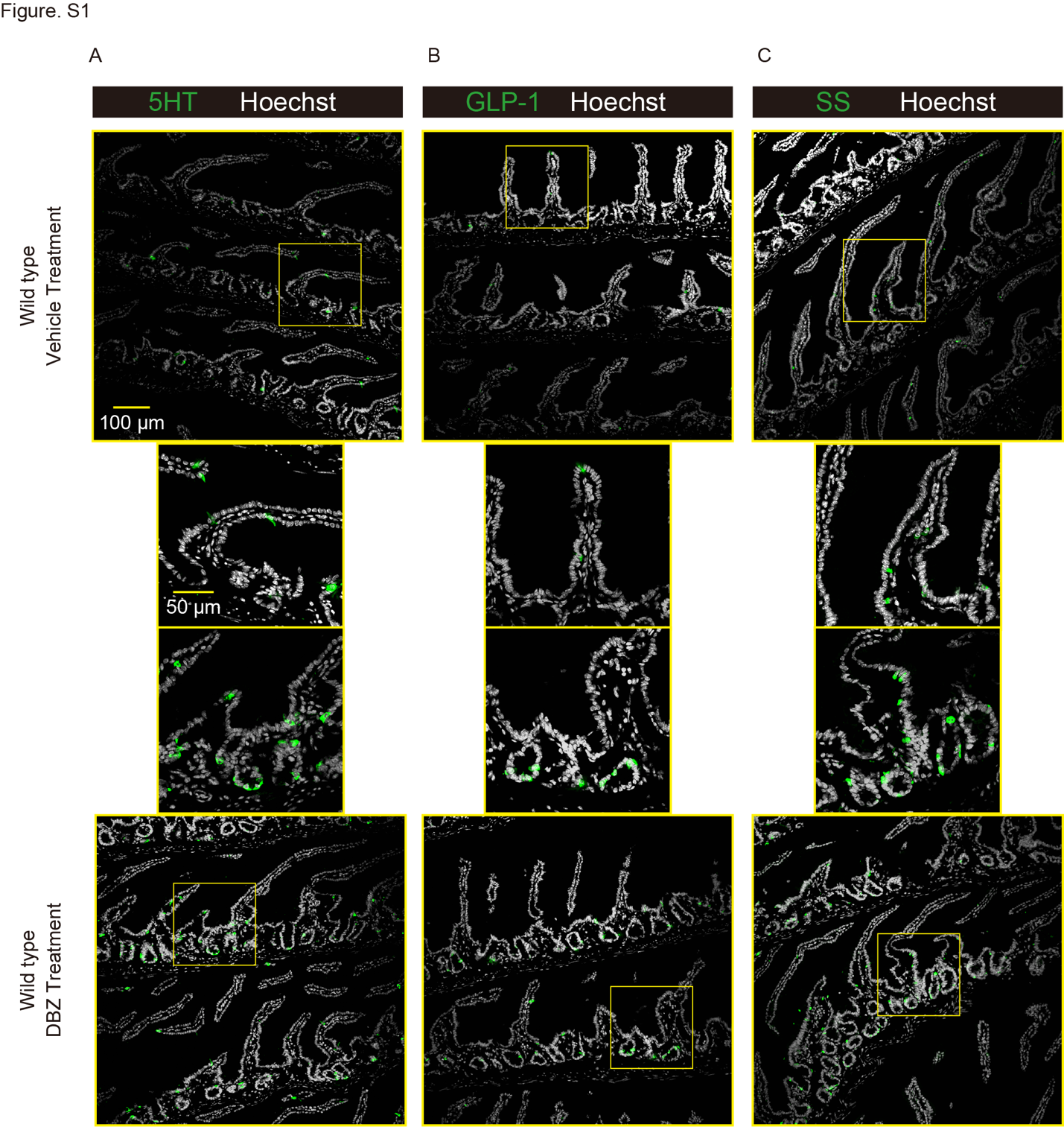
Figure S2. Induction of EECs by DBZ

Fig. S2. 5HT (A), GLP-1 (B), and SS (C) staining (green) of intestinal sections from wild-type mice 96hr after Vehicle or DBZ treatment. Scale bar = 100 μm in larger pictures, and 50 µm in magnified pictures. Hoechst counterstains nuclei.

Figure S3. Gut structure, weight, and food intake in Vil-FoxO1 knockout and wild-type control mice

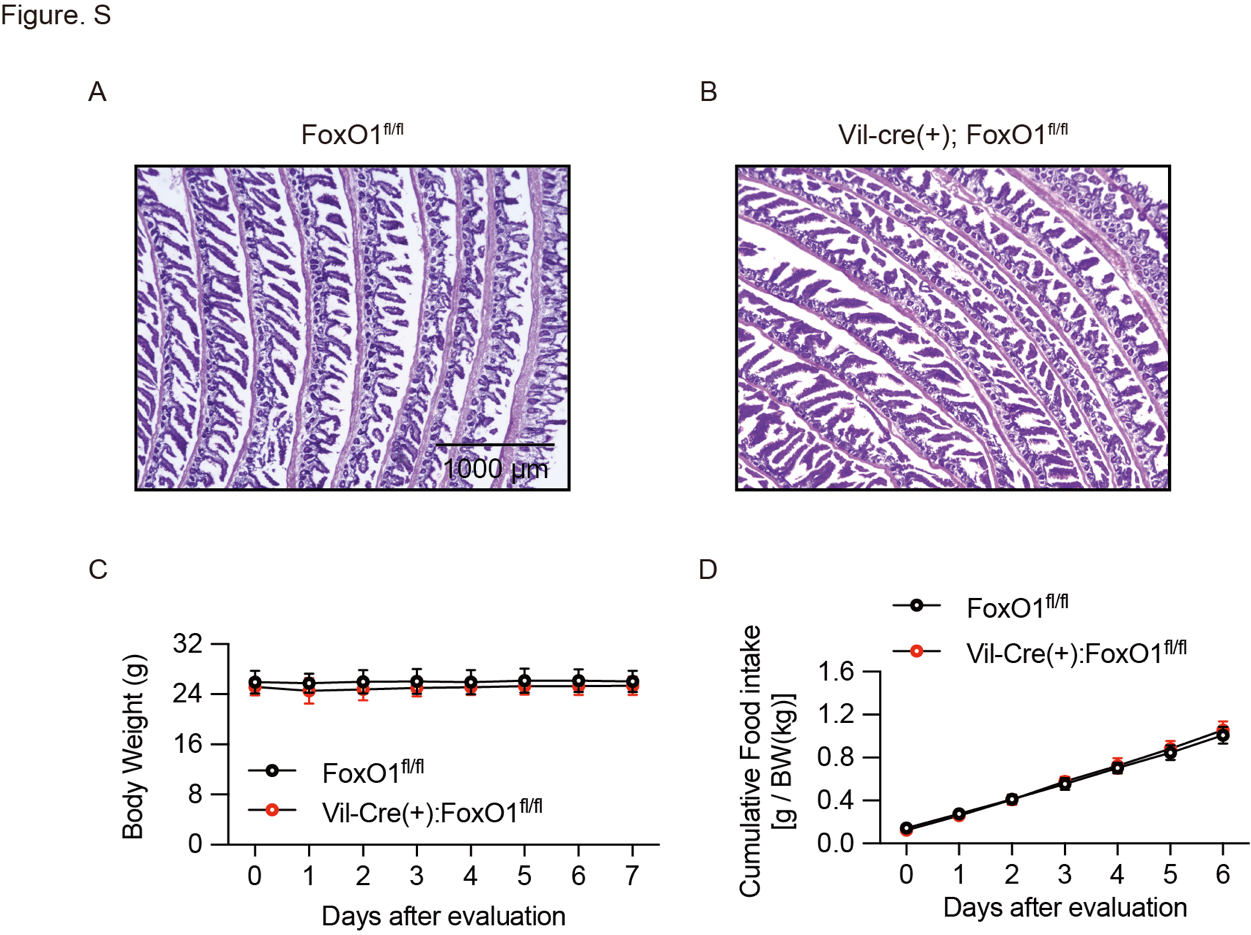

Fig. S3. (A, B) H&E of gut from FoxO1^fl/fl^ (A) and Vil-cre (+): FoxO1^fl/fl^ mice (B). (C, D) Body weight and food intake measured daily at 10 AM for a week.

Figure S4. ipGTT and OGTT in Vil-FoxO1 knockout and wild-type control mice

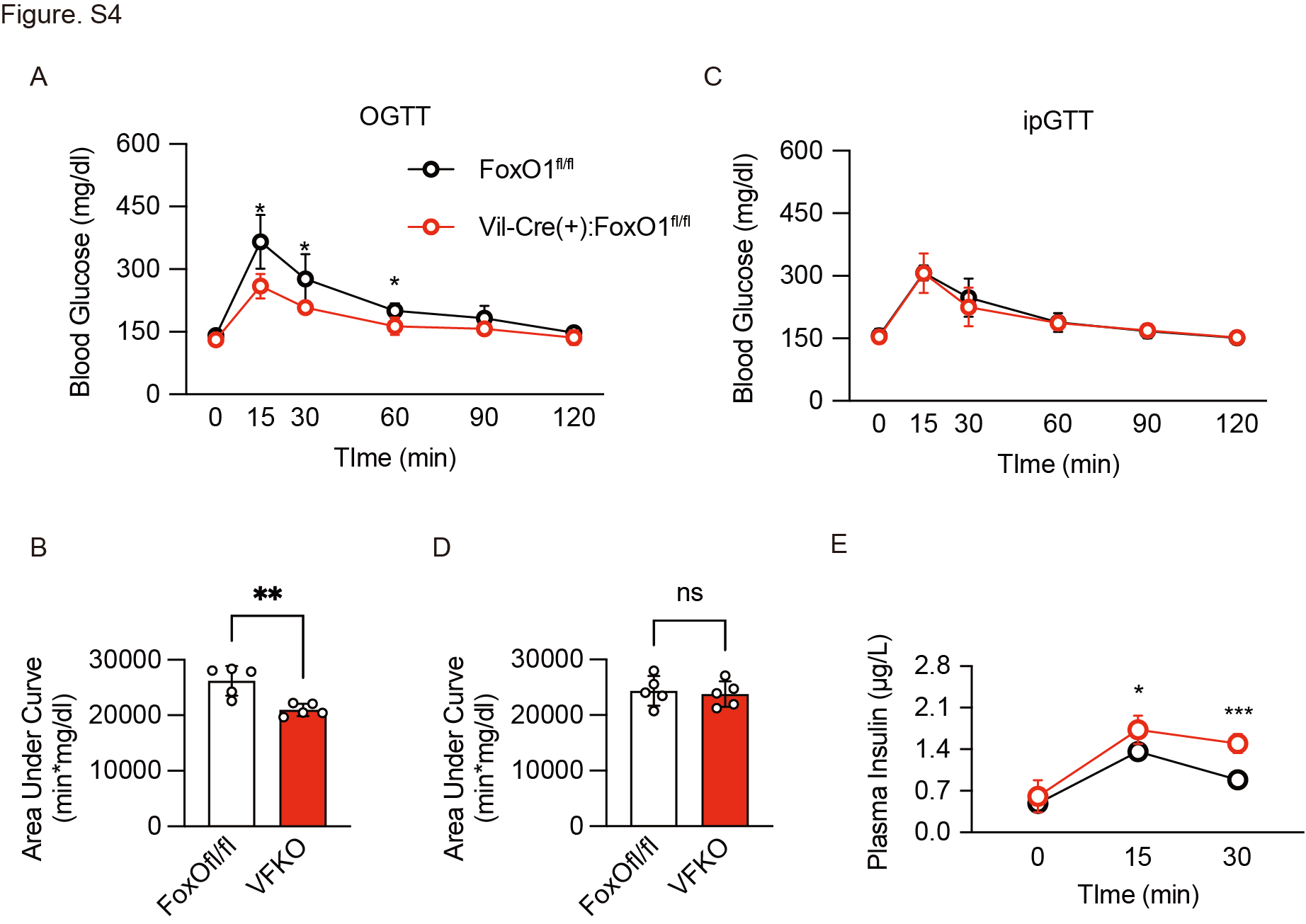

Fig. S4. (A–E) Oral (A) or intra-peritoneal (C) glucose tolerance tests (1g/kg glucose) after a 4-hr fast and area under curve of (B, D). (E) Plasma insulin levels during oral glucose tolerance tests.

Figure S5. Optimization of in vivo PF-03084014 treatment

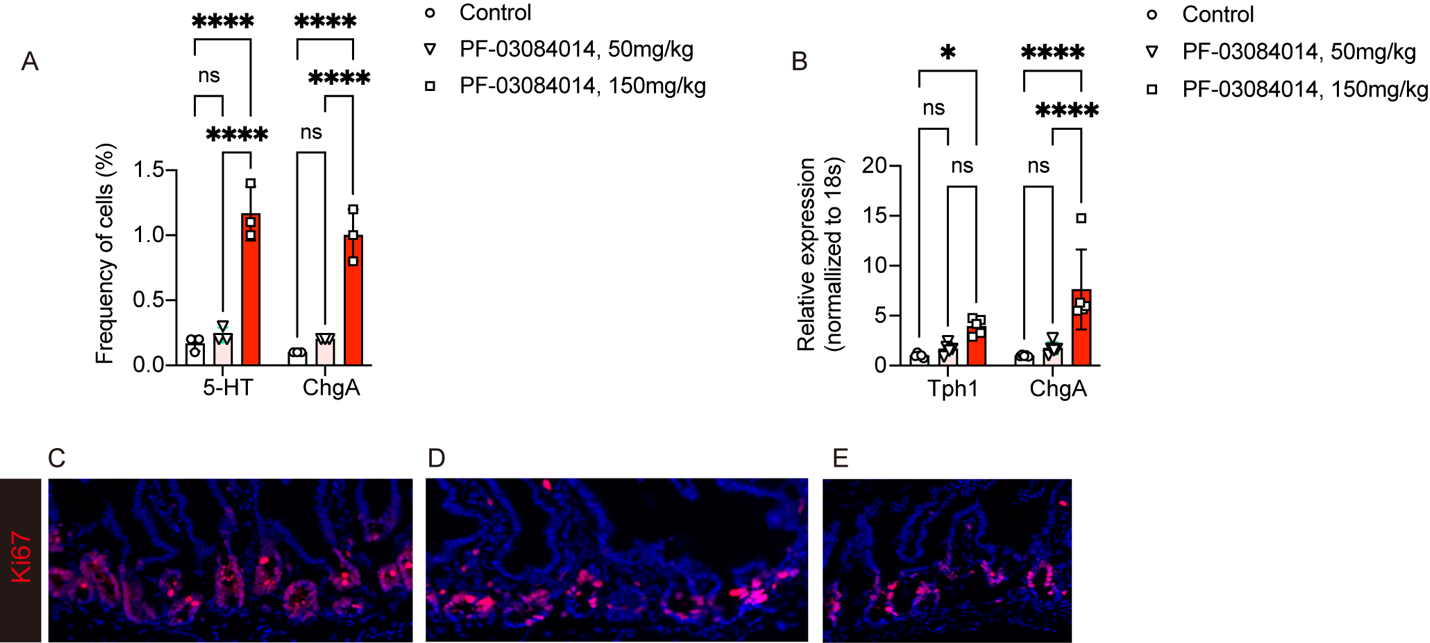

Fig. S5. (A–E) 7-8 week old C57/BL6 mice were randomized into three groups. Mice were treated with vehicle, PF-03084014 (50mg/kg/dose, PO) or PF-03084014 (150mg/kg/dose, PO) for 5 days, twice daily. (A) 5-HT and Chromogranin A cell analysis by flow cytometry after intra-cellular staining of isolated gut epithelial cells from approximately one third of the proximal end of small intestine cells (n = 3 for each group). (B) Gene expression analysis of enteroendocrine cell marker genes (Tph1, ChgA) (n = 5 for each group). (C-E) Representative images of proliferating cell marker Ki67 immunohistochemistry in tissues collected after 5 days of treatment. (C) Vehicle, (D) PF-03084014 (50mg/kg/dose), (E) PF-03084014 (150mg/kg/dose). Data are presented as means ± SD. Scale bar = 50 μm, DAPI counterstains nuclei. *= p < 0.05, ****= p < 0.0001 by two-factor ANOVA.

Figure S6. Comparison of pancreatic insulin content and quantification of GLP-1^+^, GIP^+^, and C-peptide^+^ cells in INS2^Akita/+^ mice with or without pan-intestinal FoxO1 ablation

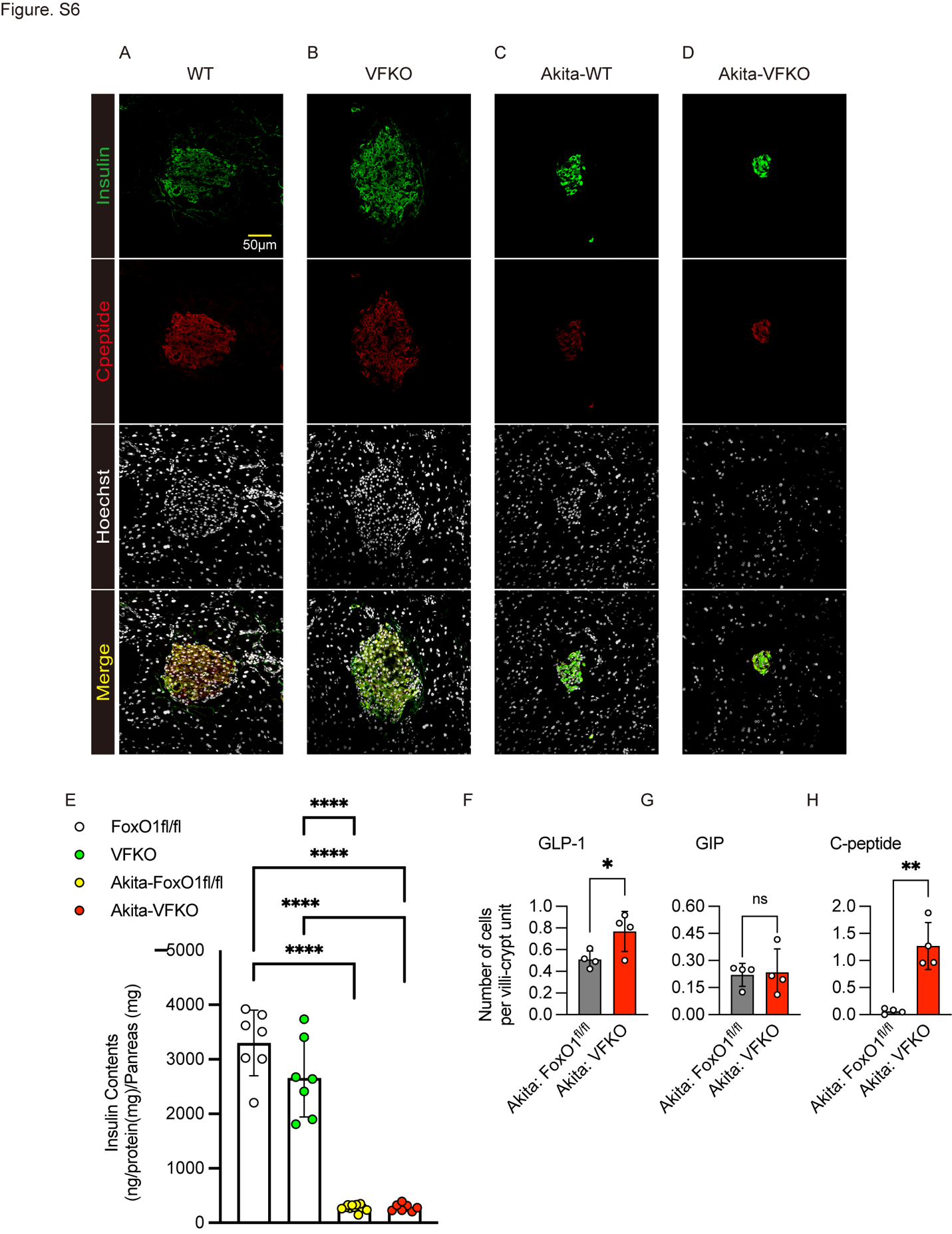

Fig. S6. (A–E) Comparison of whole pancreatic insulin content in *INS2^Akita/+^* mice with or without FoxO1^fl/fl^ and VFKO alleles. Representative islet images are stained with insulin and C-peptide (A–D). Quantification of insulin content in whole pancreas (n = 7 for each group) (E). (F–H) Quantification of GLP-1-, GIP- and C-peptide-IMMUNOREACTIVE gut cells (n = 4 for each group) in Akita-WT, INS2^Akita/+^; FoxO1^fl/fl^: Akita-VFKO, INS2^Akita/+^; Vil-Cre (+): FoxO1^fl/fl^. Data are presented as means ± SD. Scale bar = 50 μm, DAPI counterstains nuclei. *= p < 0.05, **= p < 0.01, ****= p < 0.0001 by two-factor ANOVA.

Figure S7. Effect of Cpd10 on FoxO1, 5HT, GLP-1, and C-peptide expression

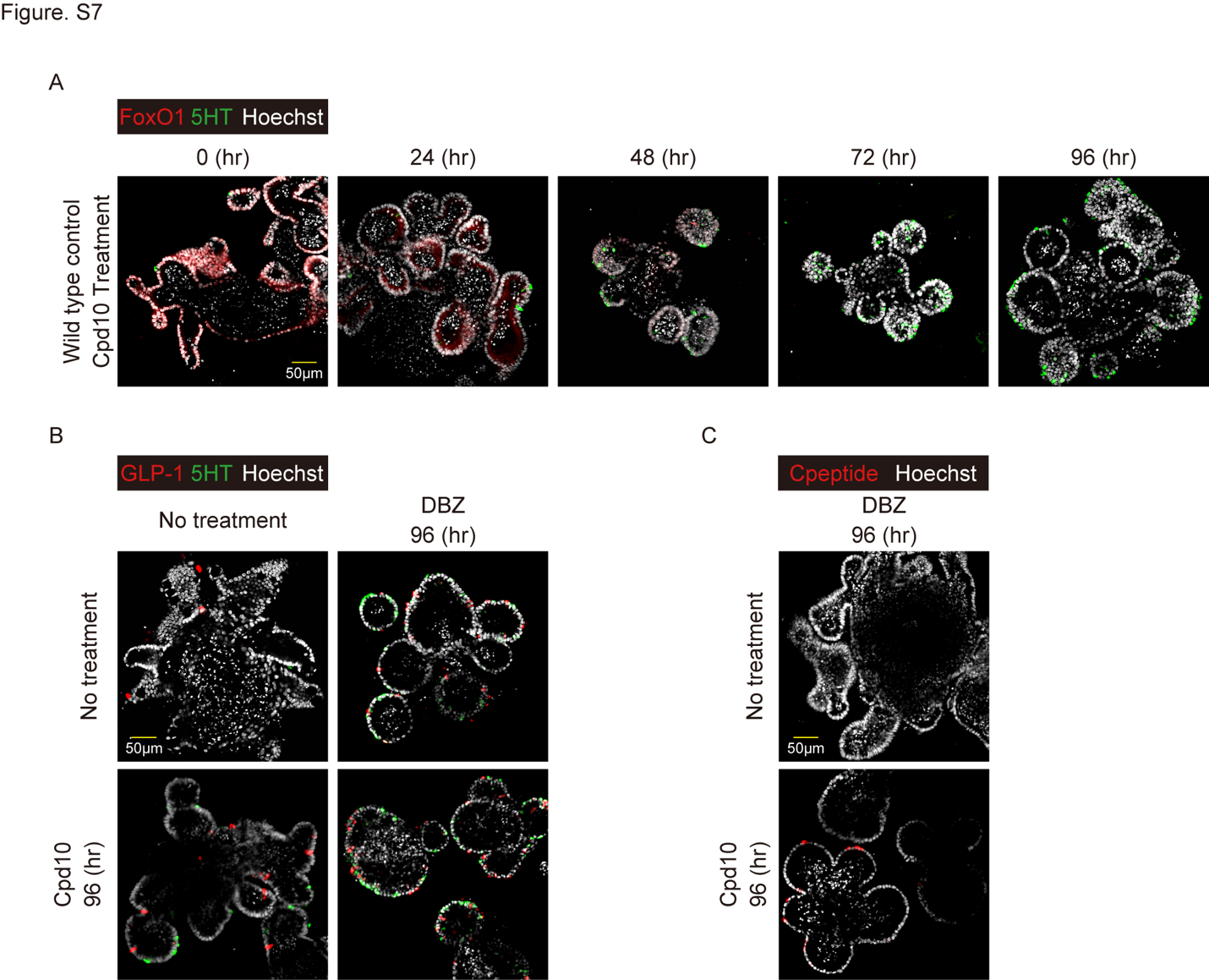

Fig. S7. (A) Time course of FoxO1 and 5HT expression of wild type mGO after Cpd10 and DBZ treatment. (B) GLP-1 immunohistochemistry of wild type mGO with or without Cpd10 treatment before and 96 hr after DBZ treatment. (C) C-peptide staining of wild type mGO 96 hr after Cpd10 and DBZ treatment. Scale bar = 50 μm. Hoechst counterstains nuclei.

Figure S8. Pharmacological effect of Cpd10 *in vivo*

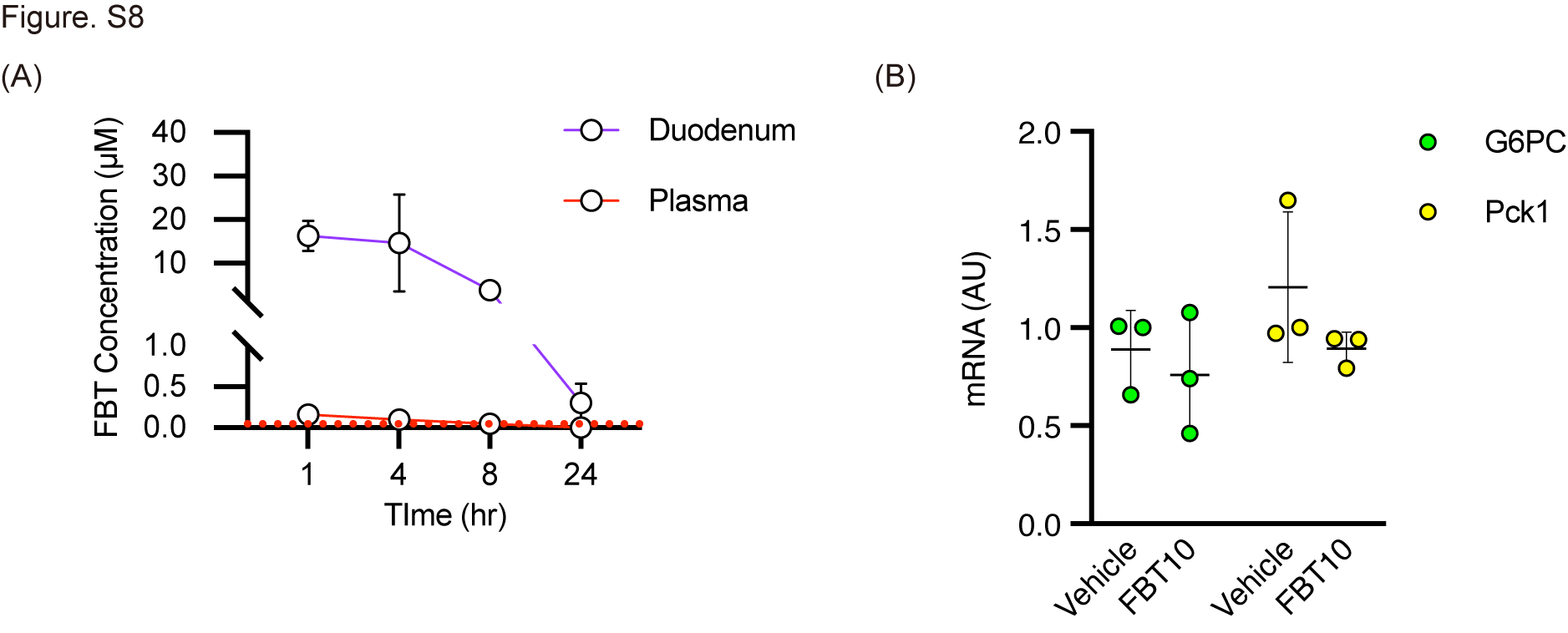

Fig. S8. (A) Gut tissue and plasma concentration of Cpd10 following a single oral dose. (B) Gluconeogenic gene expression in liver after dosing at 50mg/kg i.p injection of vehicle or Cpd10 twice daily for 4 days. G6PC: glucose-6-phosphatase, Pck1: phosphoenolpyruvate- carboxykinase. Data are presented as means ± SD. All the experiments were performed in biological triplicates.

Figure S9. Cpd10 and PF-03084014 dual treatment increases enteroendocrine cells

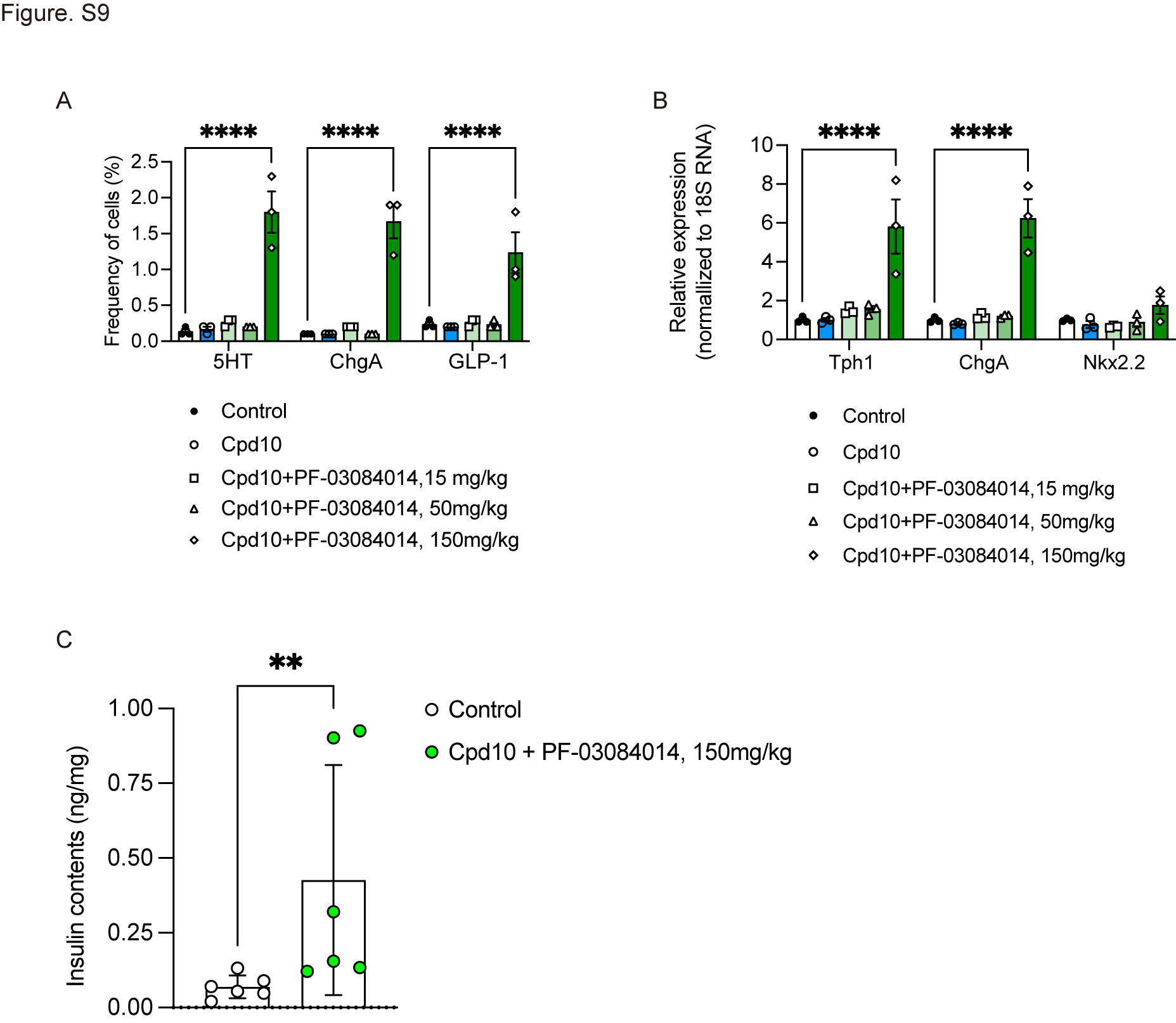

Fig. S9. (A–B) 8-9-week old Akita mice were randomized into 5 groups based on normal fed glucose levels and body weight. Mice were treated with vehicle, Cpd10 (50mg/kg/dose, PO), Cpd10 + PF-03084014 (15mg or 50mg or 150mg/kg/dose, PO) twice daily (n = 3 for each group). (A) 5-HT, Chromogranin A, and GLP-1 cell analysis by flow cytometry after intra-cellular staining of isolated gut epithelial cells from approximately one third of the proximal end of small intestine. (B) Gene expression analysis of enteroendocrine cell (Tph1, ChgA) and β-cell marker of Nkx2.2. (C) Intracellular insulin contents of gut epithelial cells from the mice treated with Vehicle or Cpd10 + PF-03084014 (150mg/kg/dose, PO) twice daily. **= p < 0.01, ****= p < 0.0001 by Mann-Whitney U test or two-factor ANOVA.

Figure S10. Representative intestinal Immunohistochemistry after 10-day single treatment with Cpd10 or PF-03084014

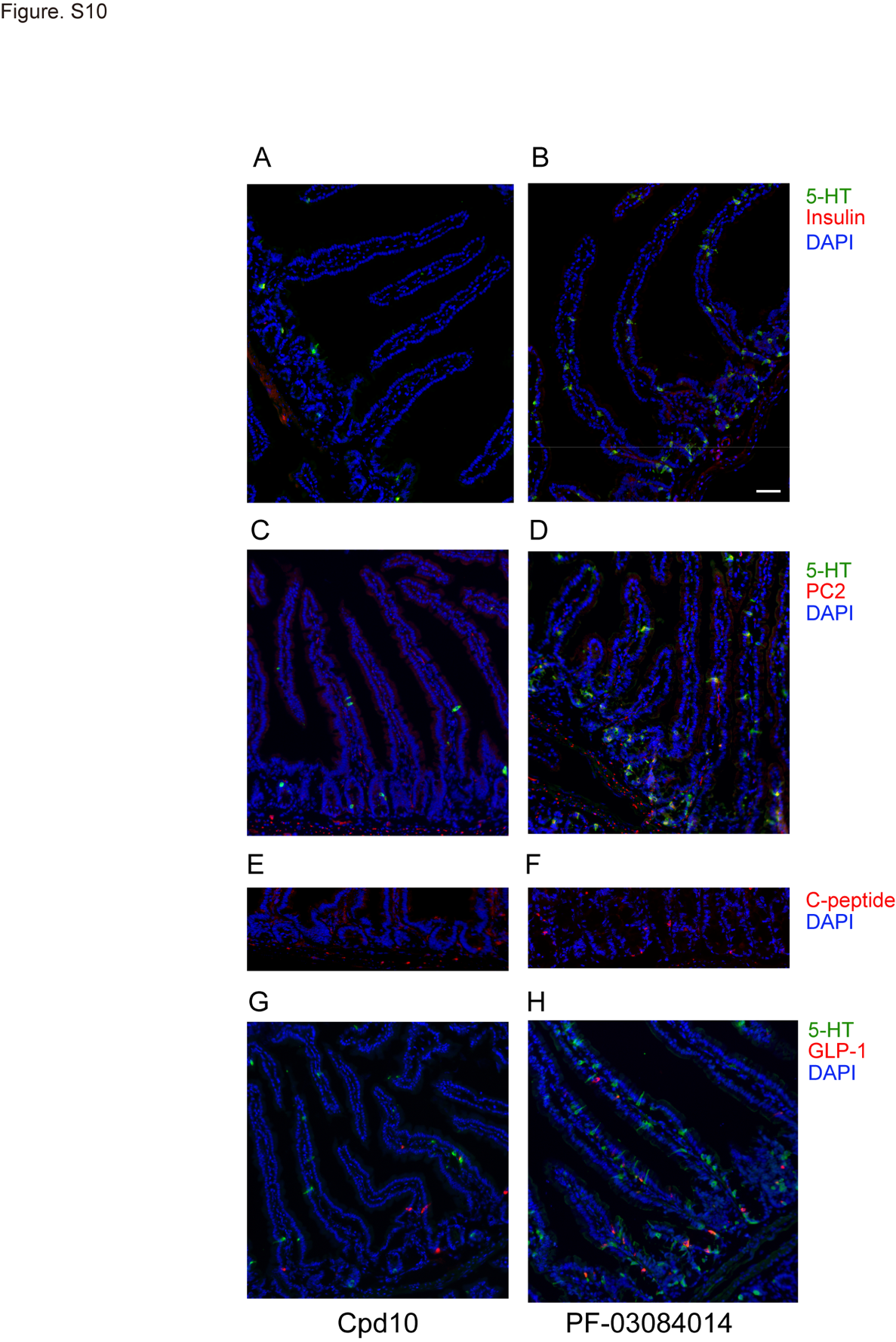

Fig. S10. (A–H) Representative intestinal immunohistochemistry after 10-day treatment. (A, B) Insulin (red) and 5-HT (green), (C, D) PC2 (red) and 5-HT (green), (E, F) C-peptide and (G, H) GLP-1 (red) and 5-HT (green) staining of Cpd10 group (left panel, A, C, E, and G) and PF-03084014-treated group (right panel, B, D, F, H). Scale bar = 50 μm

Figure S11. Images of β-cell specific markers after 10-day treatment

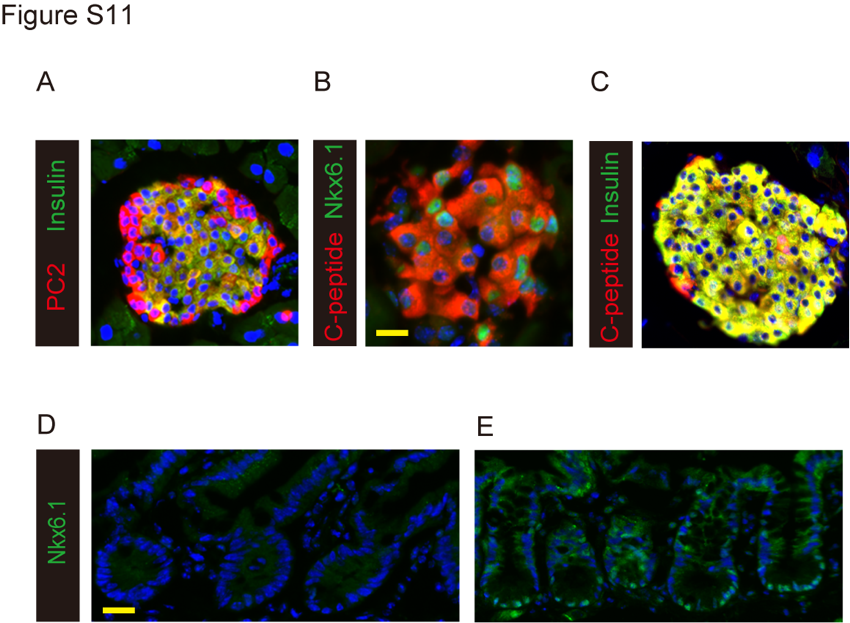

Fig. S11. (A–C) Immunohistochemistry of wild-type pancreas with beta cell markers using the same antibodies employed for gut immunohistochemistry. (A) PC2 and Insulin, (B) C-peptide and Nkx6.1, (C) C-peptide and insulin. (D–E) Representative immunohistochemistry of Nkx6.1 in the gut of 9-week-old *AKITA^Ins2 /+^* mice after 10 days treatment with Vehicle (D), or Cpd10 (50mg/kg/dose i.p) + PF-03084014 (150mg/kg/dose, PO) (E). Scale bar = 50 μm.
